# Somatic mutations impose an entropic upper bound on human lifespan

**DOI:** 10.1101/2025.11.23.689982

**Authors:** Evgeniy Efimov, Vlad Fedotov, Leonid Malaev, Ekaterina E. Khrameeva, Dmitrii Kriukov

## Abstract

Somatic mutations accumulate with age and can cause cell death, but their quantitative contribution to limiting human lifespan remains unclear. We developed an incremental modeling framework that progressively incorporates factors contributing to aging into a model of population survival dynamics, which we used to estimate lifespan limits if all aging hallmarks were eliminated except somatic mutations. Our analysis reveals fundamental asymmetry across organs: post-mitotic cells such as neurons and cardiomyocytes act as critical longevity bottlenecks, with somatic mutations reducing median lifespan from a theoretical non-aging baseline of 430 years to 169 years. In contrast, proliferating tissues like liver maintain functionality for thousands of years through cellular replacement, effectively neutralizing mutation-driven decline. Multi-organ integration predicts median lifespans of 134-170 years —approximately twice current human longevity. This substantial yet incomplete reduction indicates that somatic mutations significantly drive aging but cannot alone account for observed mortality, implying comparable contributions from other hallmarks.

## Introduction

Contemporary geroscience pursues the ultimate goal of extending human healthspan and lifespan, attracting substantial attention^1–3^. However, the very phenomenon of biological aging awaits a unified theory, and scientists worldwide engage in large-scale projects aiming to discover reliable biomarkers^4,5^, unravel aging mechanisms^6^, and develop computational models^7–9^. These findings are compiled into knowledge compendiums like the Hallmarks of Aging^10^, revealing aging as highly complex, interdependent, and multifactorial.

Some hallmarks appear reversible, and we can envision molecular pathways for drugs capable of correcting specific aging-related consequences beyond merely slowing down their accumulation. For example, telomerase overexpression reverses telomere shortening^11,12^, while senolytics target senescent cells^13,14^, albeit actual longevity therapies are yet to appear. Other aging phenomena represent pure information loss (or entropy increase)^15–18^. One prominent example is somatic mutations — DNA alterations that occur throughout lifetime in non-germline cells^19,20^. Although considered part of genomic instability^10,21^ (Fig. 1a,b), their impact on aging and life expectancy has been debated^22,23^. Mounting evidence suggests somatic mutations are not mere clock-like changes^24^, but drastically affect health by altering epigenetic landscapes^25^, inducing within-tissue genome mosaicism, generating pathogenic clones, and enabling age-related diseases^26–31^.

**Figure 1.**
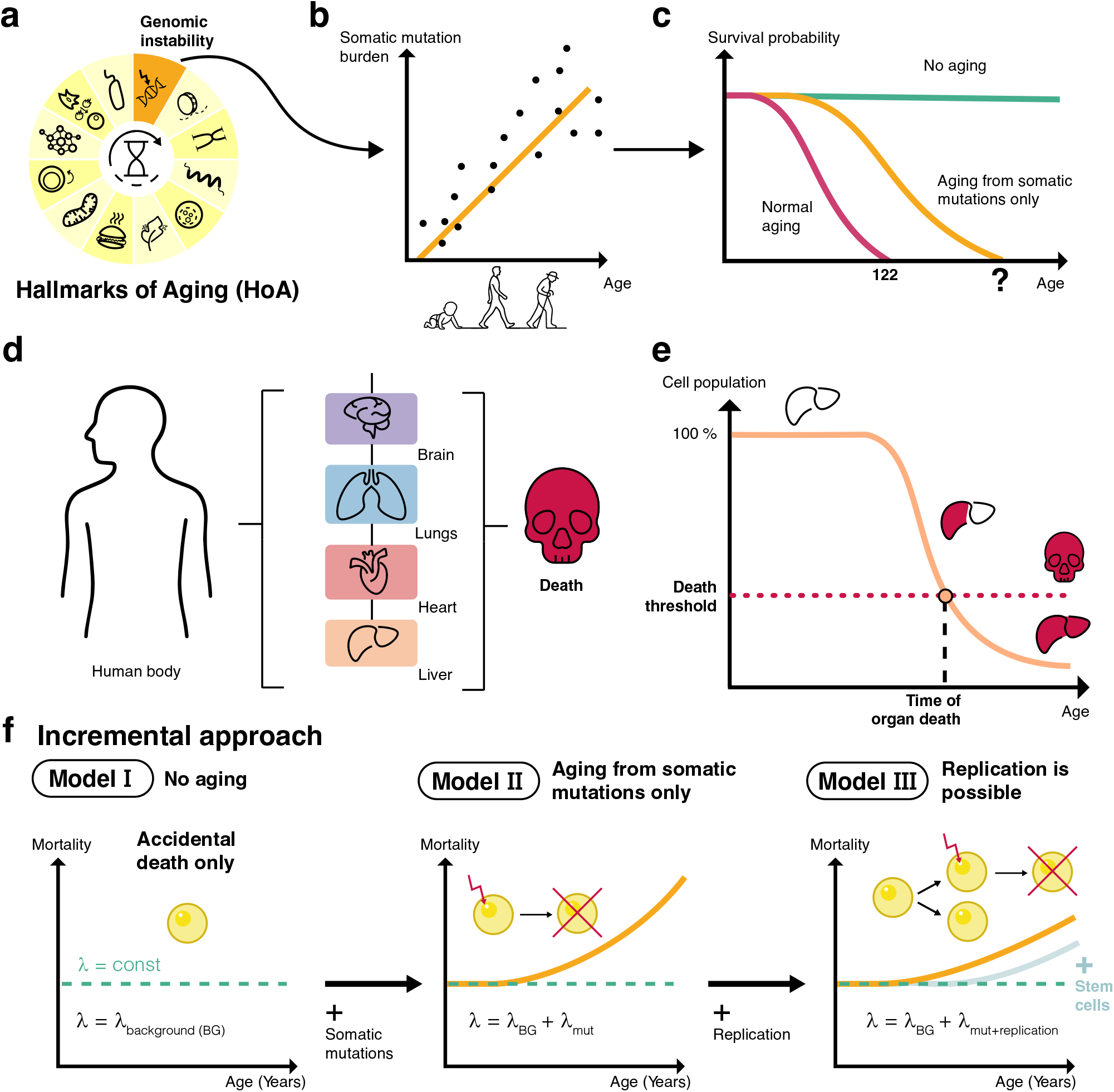
Hazard rates and lifespans estimated for different ages in different populations. According to Model I, maximum and median lifespans are defined as ages at which the survival probability drops to 1*/N* (where *N* is the global human population) and to 0.5 (or 50%), respectively. The lifespans calculated for the 30-year mortality are highlighted with black frames and larger font size.

Like any mutation, the somatic ones arise from infidelity in DNA damage repair, replication, or mitosis^32^. While natural selection strives to preserve DNA integrity, perfecting its maintenance is too costly in resources and time alternatively spent on transcription and proliferation^33^, so species tolerate mutation rates allowing their survival^34^. The disposable soma theory^35^ implies that germline genetic information must be protected by all means, while somatic DNA is not inherited, so it needs less fidelity^32,36^ (experimentally confirmed by higher somatic versus germline mutation rates^19,20,37^). Accumulated somatic damage causes cell function to decline, leading to diseases and organismal aging^38^. Specific recurrent nucleotide alterations (mutational signatures) may confer cell survival benefits, causing clones harboring these signatures to proliferate faster and spread mutations widely in mitotically active tissues through clonal expansion events (e.g., clonal hematopoiesis), further accelerating aging^27,30,39^.

While there are approaches that might eventually mitigate somatic mutations (including elimination of high-burden cells^32^, whole-genome editing^32,40^, or harnessing revertant mosaicism^41,42^), they remain extremely far from clinical application. Consequently, hypothetical anti-aging therapies might only reverse conditionally reversible hallmarks while being fundamentally unable to affect entropic phenomena like somatic mutations. To date, no attempts have been made to estimate how long a future human, having undergone a treatment against reversible aging, could possibly live (Fig. 1c). What would be the maximum effect of radical life extension^43,44^ excluding extravagant interventions like massive cell replacement^45,46^ or transferring consciousness onto a silicon chip^47–49^? In other words, what is all this strife for?

This work aims to estimate how long humans would live if we cured most aging processes except somatic mutations (Fig. 1c). Given accumulated sequencing data, relative simplicity of mutation mechanisms^50^, and, on the other hand, virtual irreversibility of somatic mutation burden, focusing on this phenomenon provides a plausible assessment of human lifespan’s upper bound.

Previous models evaluating the relationship between somatic mutation accumulation and lifespans found that mutation rates are inversely proportional to mammalian species lifespan^51,52^. While intriguing, this oversimplifies the dynamics of mutation-driven aging without providing frameworks for theoretical investigations. Other dynamical systems assessing cell population dynamics focus mainly on stem cells^53^ and are better suited for studying cancer development^54,55^ rather than normal aging.

Estimating maximum possible lifespan while theories of aging remain immature is challenging and requires its own novel methodology. Comprehensive estimation would require complex models of somatic mutation dynamics across all tissues and organs plus a host of public data on mutation rates and stem cell differentiation across ages, tissues, and individuals. However, guided by reliability theory, we instead represent the human body as a system of conditionally-independent components^56,57^ (Fig. 1d), configured in parallel (redundantly) or in series (non-redundantly). This model predicts that if any unique component (e.g., brain, heart, liver) or all instances of parallel components (both lungs or kidneys) reach critical malfunction — due to cell death, in the simplest scenario (Fig. 1e) — the whole system dies, as do the real organisms unless organ function is restored artificially. We can thus interrogate each essential organ individually, focusing on those which are well-studied in terms of somatic mutation burden.

Another technique routinely used in mathematical and physical reasoning involves evaluating lower and upper bounds of given parameters. For terminally differentiated, non-dividing cells like neurons, stochastic death from deleterious mutations would provide an upper lifespan bound, considering brain failure equivalent to organism death. When modeling other tissues, incorporating cell division and differentiation would markedly improve the estimates.

In our work, we propose an incremental approach to modeling aging, which involves incorporating aging factors progressively to increase model complexity when necessary (Fig. 1f). We developed dynamical system models describing particular critical organs that deteriorates under somatic mutation burden, with model complexity adjusted being adjusted based on tissue replication capabilities. These models apply reliability theory to biological systems and employ upper/lower bound estimates for variables lacking solid experimental evidence. By refining and generalizing the models — first per organ, then within interconnected critical blocks (Fig. 1d) — we enhance their capacity to reflect biological reality and estimate lifespans. Applying this framework, we leverage data from brain, heart, liver, and lungs, offering an in-depth view of organ-specific cell population dynamics under somatic mutation pressure, allowing us to estimate the limits to human lifespan when aging is driven purely by somatic mutagenesis. We thus present a robust framework for evaluating the impact of somatic mutations on aging (Fig. 1c–e) and propose a novel approach to modeling aging, which could potentially enable ranking of fundamental aging mechanisms by their contribution to organismal aging, and determining which should be prioritized in our efforts to counteract them.

## Results

### Designing an incremental approach for modeling aging under somatic mutations

Before modeling aging, we must define it precisely. Despite numerous theories, no consensus exists^58,59^, except that aging ultimately manifests as a demographic phenomenon through population-level survival curves. Here, we adopt a working definition of aging as the age-dependent increase in mortality risk (also known as hazard rate). Modeling large-scale population aging while accounting for all mechanistic biological factors presents a formidable challenge. Existing models capture only specific molecular aspects (e.g., senescent cell accumulation^60^) or omit detailed mechanisms in favor of phenomenological perspectives, treating organisms as systems that accumulate abstract damage^17,61,62^.

To combine biological factors with population-level survival, we designed an incremental strategy. At each step, we introduce an elementary survival-contributing factor, progressively refining the model. Our baseline (Model I) describes a non-aging population where all individuals have constant mortality rates. Next, we view every individual as a system of critical elements (organs), a failure of each causing instant death. Organs fail when a certain percentage of their cells die. We focus solely on cell death from somatic mutations, assuming all other death causes are incorporated into background mortality from Model I and do not increase with age. Model II considers background mortality plus cell loss from somatic mutagenesis. Model III adds replication for actively proliferating tissues, in three variants. Model IIIA includes somatic cells with replication limited by proliferative potential (Hayflick limit^63,64^). Model IIIB includes somatic cells plus adult stem cells which replenish somatic pools via differentiation, possess unlimited proliferative potential, but die from mutations as well. Model IIIC describes stem-like cells with more prolonged but still limited proliferation that leads to both self-replenishment and differentiation. Accounting for variability in maximum organ capacity and cell death/replication rates yields varying cell population trajectories per organism, incorporating cell-level uncertainty when calculating population-level survival curves.

Finally, we integrate individual aging organs into a single system with the reliability theory framework, yielding combined survival trajectories that unite background and organ-specific mortality due to somatic mutations.

### Model I: Baseline model of a non-aging population yields an enormous theoretical maximum lifespan

The first step of our framework establishes a baseline model to describe mortality dynamics in the complete absence of aging. Technically, a non-aging population is assumed to have constant mortality rate throughout lifespan, allowing to calculate its survival for Model I (Fig. 2a) as:

**Figure 2.**
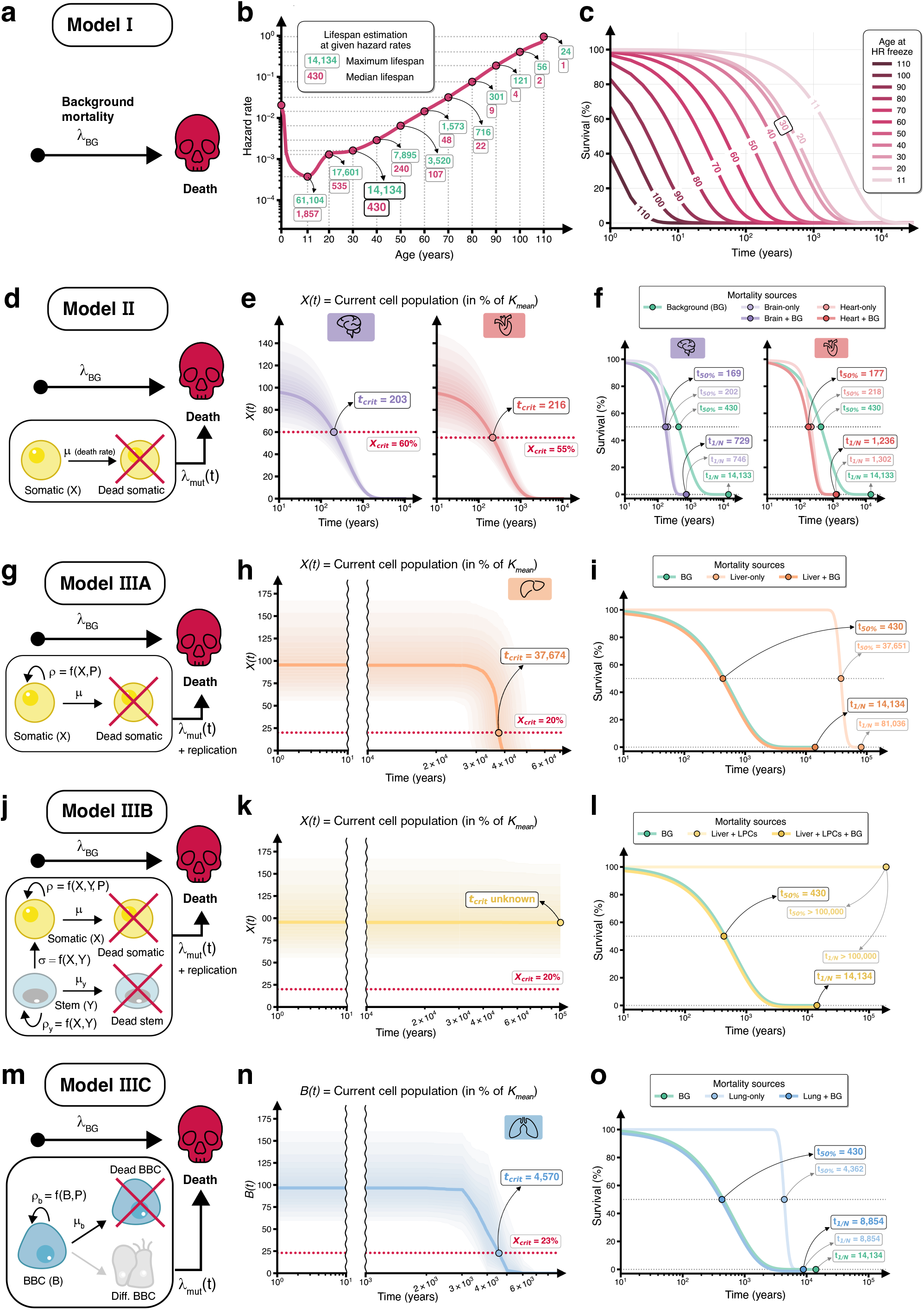
Estimation of SNV and indel accumulation rates *µ*_0_ across human lifespan in cell types included in the study.

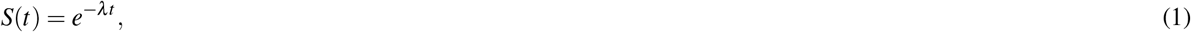

where *λ* is constant mortality rate from demographic data fixed at a specific age, *t* is time elapsed since birth, and *S*(*t*) is population survival function. The Gompertz law^65^ of exponentially increasing mortality degenerates into constant mortality. But how to determine this constant *λ* ?

From empirical life tables (http://www.lifetable.de), we can evaluate mortality rates at different ages and solve equation (1) for *t* (see Methods). We can thus obtain the lifespan of a population with “frozen” mortality *λ* as the value of *t* at any *S*(*t*) (percentage remaining from the initial population), assuming its initial size as *N* = 8 billion individuals — approximately current human population^66^. We define median survival as time until half the initial population survived and maximum survival as time until one person from *N* survived, yielding:

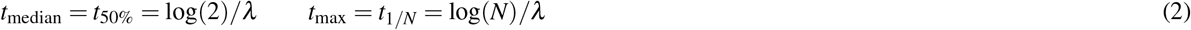

Naturally, mortality rates at different ages and survival curves vary by population. We used life tables to estimate median and maximum lifespans for nine countries (Extended Data Fig. 1) and focused on the US data (Fig. 2b,c) for downstream modeling. Importantly, non-aging populations are not immortal: death causes continue acting on individuals, inevitably leading to eventual extinction, which is reflected by survival curves for mortality rates fixed at every ten years: all of them eventually fall to nearly zero (Fig. 2c).

All nine studied populations display J-shaped curves of mortality rate, with their minima lying between 5–15 years depending on the country, followed by steep exponential rise appearing as linear growth in the logarithmic scale, with some populations (US, China, Germany, Israel) exhibiting intermediate mortality plateaus between ages 20–30 (Extended Data Fig. 1).

We calculated median and maximum lifespans for several age points of mortality “freeze” (Fig. 2b, Extended Data Fig. 1). For the US population, mortality fixed at age 20 yields median lifespan of 535 years and maximum of 17,601 years. Though extraordinary, these numbers follow directly from our framework. Conversely, fixing mortality at the 110-year-old supercentenarian levels yields median remaining lifespan of only 1 year — consistent with intuition. The maximum remaining lifespan is 24 years, reflecting probabilistic survival of at least one individual among 8 billion under such elevated mortality risks, which aligns with the reflections of the late Mikhail Blagosklonny who noted that “A mere application of standard medical care to centenarians, as rigorously as to younger adults, would probably extend lifespan beyond 122, even without the need of a scientific breakthrough.”^67^

The childhood minimum mortality point (11 years in the US dataset) corresponds to highest predicted lifespans, both median (1,857 years) and maximum (61,104 years) — a striking illustration of how age-dependent differences in mortality profoundly affect projected lifespan in our non-aging model.

The choice of baseline mortality risk links to the question: “At what age does aging begin?” Perspectives on this issue vary widely^58,59^ — from aging starting during embryogenesis^68^ to the arguments that it begins only when body growth ceases, consistent with disposable soma theory^35,69^. Mortality risk curves (Fig. 2b, Extended Data Fig. 1) reveal that the exponential mortality increase typically emerges around age 30 — a threshold coinciding with the commonly recognized onset of middle age. We adopt the age 30 mortality rate as our baseline, which corresponds to median and maximum remaining lifespans of 430 and 14,134 years. This choice reflects the mortality risk of a typical early adult characterized by good health, low incidence of age-related pathologies, and full societal engagement. At this age, individuals are generally active, employed, and routinely exposed to diverse extrinsic hazards including car accidents, drug abuse, occupational hazards, infectious diseases, etc., thus embodying composite baseline mortality risks without the influence of aging.

Consequently, all further analyses assume that mortality rate is fixed at the levels of a modern 30-year-old person and referred to as background mortality (*λ*_30 y.o._ = *λ*_BG_). Building upon this assumption, we incrementally incorporate additional factors related to somatic mutagenesis and cell proliferation that contribute to organismal survival, progressively enhancing model realism and explanatory capacity.

### Model II. Somatic mutations in post-mitotic tissues drive substantial mortality increase but cannot fully account for observed aging

Building upon the baseline non-aging model, we next introduced somatic mutagenesis in non-dividing cells, focusing on neurons and cardiomyocytes as representative post-mitotic cell types (Fig. 2d-f). Cell loss dynamics driven by deleterious mutations can be described by an ordinary differential equation (Model II):

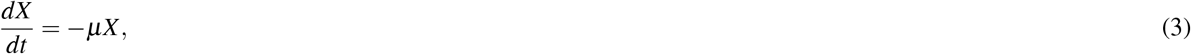

where *X* is the current number of cells at risk in an organ, *dX* is the number of cells lost over time *dt*, and *µ* = *µ*_0_ · *p*_lethal_ is death probability per cell, which equals to the product of: *µ*_0_ — somatic mutation accumulation rate measured as number of mutations per cell per year, and *p*_lethal_ — the probability of a single mutation to be lethal for a cell. Both *µ*_0_ and *p*_lethal_ can be estimated separately for every mutation type — we focused on single nucleotide variations (SNVs) and insertions-deletions (indels) — and then summed up for a combined estimate of *µ* (see Methods).

Equation (3) for Model II corresponds to an exponential decay solution, *X* (*t*) = *Ke*^−*µt*^, with decay rate governed by the *µ* coefficient, and starting from organ cell capacity *X* (*t* = 0) = *X*_max_ = *K*.

From the literature describing how somatic mutation burden accumulates with age across organs^70–73^, we estimated *µ* for two essential terminally differentiated cell types: cortical neurons (*µ* ≈ 2.44 · 10^−3^ lethal mutations per cell per year) and cardiomyocytes (*µ* ≈ 2.61 · 10^−3^) (see Methods; Extended Data Fig. 2, Table 3). We assumed that organ failure occurs once the number of functional cells falls below a critical fraction determined from empirical physiological thresholds (see Methods).

To translate cell-level dynamics into organism-level survival, we determined when each organ reaches its critical failure threshold based on the exponential cell loss from equation (3). Accounting for variability in both decay rate *µ* and initial organ capacity *K*, we computed the times to reach organ failure (Fig. 2e). We then combined this somatic mutation-driven mortality with baseline mortality from Model I by summing their respective hazard rates *λ*_mut_ (organ-specific) and *λ*_BG_, assuming independence between mortality sources that is equivalent to multiplying their survival functions (see Methods):

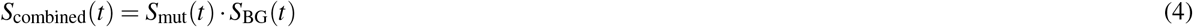

Combining baseline mortality (fixed at age 30) with somatic mutation-driven cell death yields a population survival curve with median lifespan of 169 years and maximum of 729 years for neuron-driven aging, and 177 and 1,236 years, respectively, for cardiomyocyte-driven aging (Fig. 2f). Compared to the non-aging baseline (median: 430 years, maximum: 14,134 years), this represents a substantial reduction driven by a single elementary aging mechanism. However, the resulting maximum lifespans remain markedly longer than the observed maximum of 122 years, while the resulting median lifespans are twice as long as median life expectancies in developed countries, indicating that somatic mutations in post-mitotic tissues, while contributing substantially, cannot alone explain observed human mortality patterns.

### Model III: Cellular replication in proliferating tissues provides strong protection against somatic mutation-driven aging

The previous model applies only to post-mitotic tissues that cannot replace lost cells. However, most human tissues comprise actively proliferating cells capable of compensating for cell loss through replication. To determine whether cellular replication can mitigate somatic mutation-driven aging, we extended our model to include both dividing somatic cells with finite replicative potential and a pool of stem cells that self-renew and differentiate (see Methods):

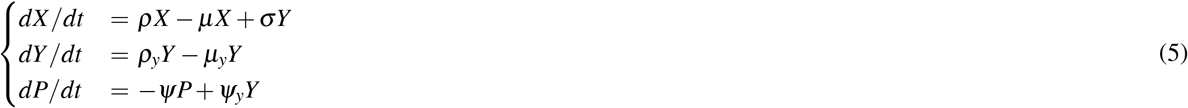

where *µ* and *µ*_*y*_ represent cell death rates for somatic and stem populations, while *ρ* and *ρ*_*y*_ represent their respective replication rates. Somatic cells possess limited proliferative potential *P* (Hayflick limit), which depletes with each division at rate *ψ*, but can be replenished at rate *ψ*_*y*_ through stem cell differentiation, which adds cells to the somatic pool at rate *σ*. All these rates (except the death rates) are functions that depend on current somatic/stem cell populations (see Methods).

Although the resulting system of differential equations lacks a closed-form analytical solution, it can be solved numerically. Using empirical data, we estimated the required parameters (see Methods; Extended Data Figs. 2, 7, 8, 9, 11, and 12) for two representative proliferating tissues: liver (hepatocytes and liver progenitor cells, LPCs) and respiratory epithelium (bronchial basal cells, BBCs). We implemented three variants of this model.

#### Model IIIA: hepatocytes, finite replicative capacity, no stem cells

Simulation results for Model IIIA (*Y* = 0; Fig. 2g-i) reveal that even without the LPC support, hepatocyte replicative capacity provides robust protection against somatic mutation-driven cell loss. Median time to reach the critical threshold extends to 37,669 years, far exceeding failure times in post-mitotic organs (729 years for neurons, 1,236 years for cardiomyocytes). Consequently, liver failure contributes negligibly to population-level mortality: the median and maximum lifespans remain virtually unchanged at 430 and 14,134 years (Fig. 2h,i), matching the non-aging baseline.

#### Model IIIB: hepatocytes + liver progenitor cells

Model IIIB (calculated using full equation (5); Fig. 2j-l) incorporates LPC-mediated hepatocyte replenishment and yields even more striking results. None of the simulated trajectories reached organ failure within 100,000 years, yielding 100% survival (Fig. 2k,l). These findings indicate that when supported by stem cells, liver exhibits exceptional resilience to somatic mutation-driven damage and does not constitute a lifespan-limiting bottleneck. Hence, liver mortality can be safely neglected within our modeling framework.

#### Model IIIC: respiratory basal cells

Model IIIC for respiratory basal epithelial cells (see Methods; Fig. 2m-o) reveals somewhat distinct aging patterns compared to Models IIIA and IIIB. Despite finite replicative capacity and continuous loss through differentiation, basal cells initially maintain stable population dynamics: cell population trajectories remain nearly horizontal for thousands of years before declining linearly as cells exhaust their proliferative potential and succumb to accumulated mutations. Median cell population trajectory crosses the critical threshold at 4,570 years (Fig. 2n). This delayed decline manifests in the organism-level survival curve as prolonged stability at 100% survival, followed by a rapid drop, yielding median lifespan of 4,362 years and maximum of 8,854 years (Fig. 2o). When combined with baseline mortality, the resulting population survival curve closely resembles the non-aging baseline throughout most of the lifespan, with median survival remaining at 430 years. However, unlike Models IIIA and IIIB, maximum lifespan becomes reduced from the baseline to 8,854 years. These results indicate that while respiratory epithelium does not limit median lifespan, it may eventually impose an upper bound on maximum survival, suggesting that even tissues with substantial regenerative capacity can become lifespan-limiting factors at extreme ages.

Collectively, these findings demonstrate fundamental asymmetry in how somatic mutations affect aging across tissue types. While somatic mutations drive substantial mortality increases in post-mitotic tissues, incorporating cellular replication effectively neutralizes this effect in studied proliferating tissues (Extended Data Figs. 4 and 5). Across all three replication-based models (IIIA, IIIB, and IIIC), organ failure times extend far beyond the lifespans observed in normal aging and projected for non-aging, rendering the individual contribution of somatic mutations negligible in actively proliferating tissues.

### Multi-organ model reveals theoretical lifespan limits under pure somatic mutation-driven aging

We now integrate these organ-specific models into a multi-organ framework inspired by the reliability theory^56^. An organism is represented as a series system of organs, surviving only if all critical organs remain functional, i.e., failure of any single organ causes death (Fig. 3a). This series configuration also includes an additional element representing background mortality *λ*_BG_, as defined in Model I. Assuming independence among aging processes in individual organs, the organismal survival curve can be expressed as the product of each organ’s survival functions and baseline survival (Fig. 3a).

**Figure 3.**
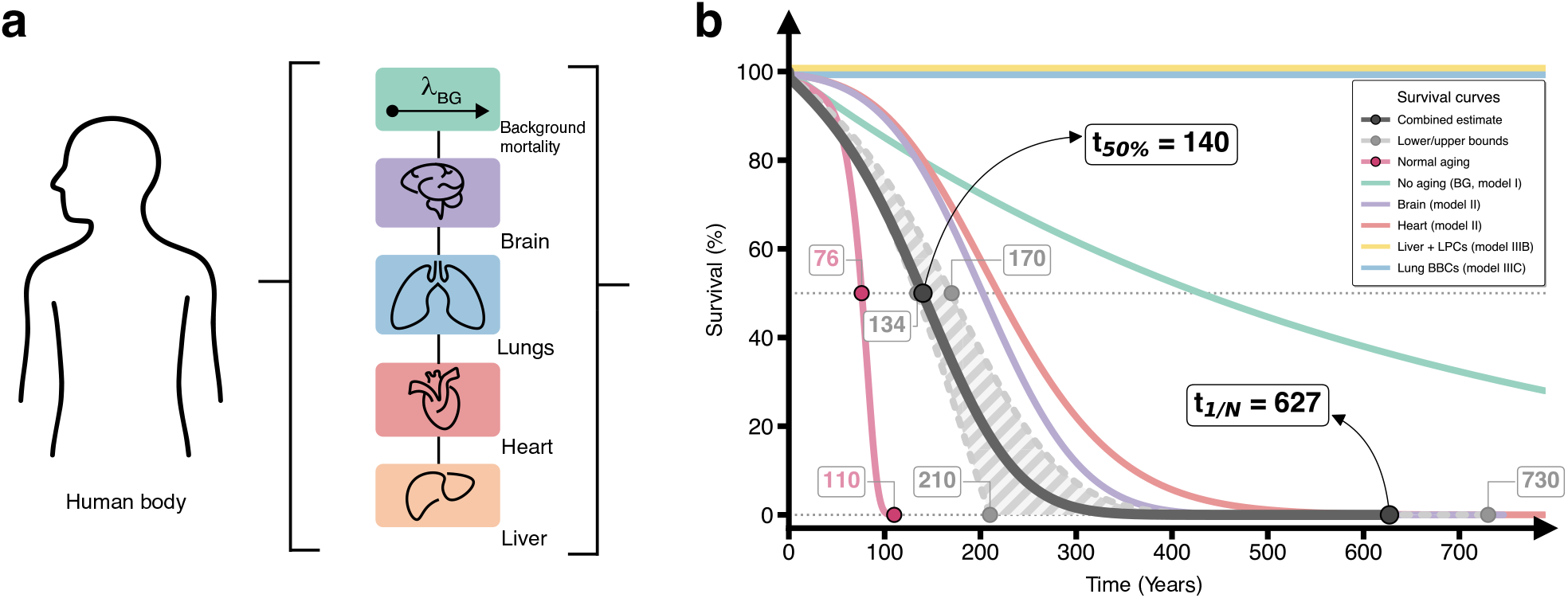
Survival curves obtained by applying Model II for all organs included in the study. Horizontal dotted gray lines correspond to survival probabilities of 0.5 and 1*/N*; median and maximum lifespans are annotated for organ-only and combined survival curves as *t*_50%_ and *t*_1*/N*_, respectively.

The resulting composite survival curve describes a hypothetical human population aging solely through somatic mutations in four critical cell types (neurons, cardiomyocytes, hepatocytes, and respiratory basal epithelial cells), with background mortality fixed at 30-year-old levels. According to this model (Fig. 3b), estimated median lifespan is 140 years, corresponding to 184% of the empirically observed median human lifespan of 76 years. Maximum lifespan reaches 627 years, or 514% of the empirically observed maximum of 122 years.

The independence assumption represents a strong simplification of biological reality, because organs do not age in isolation. To quantify uncertainty arising from potential inter-organ dependencies, we applied Fréchet inequalities^74^ (see Methods). These inequalities establish upper and lower bounds for the joint survival probability. The upper bound assumes perfect positive dependence — all organs age in perfect synchrony, as if their health states were perfectly correlated. In this scenario, organismal survival is limited by whichever organ fails first. The lower bound assumes perfect negative dependence — one organ’s integrity maintained at another’s expense. While neither extreme is biologically realistic, they provide conservative boundaries that quantify maximal possible deviations from independence.

Under these bounds, median lifespan ranges from 134 years (perfect negative dependence) to 170 years (perfect positive dependence) (Fig. 3b). Maximum lifespan shows greater sensitivity to inter-organ dependency, spanning 210–730 years across the same range of bounding assumptions. This broad interval for maximum lifespan reflects compounding effects of dependency assumptions when extrapolating to extreme population tails.

## Discussion

Motivation for this study stems from a deceptively simple yet profoundly consequential question: *What if we were to eliminate all reversible mechanisms of aging, leaving only irreversible, entropy-driven processes such as somatic mutations? How long could a human then live?* To address this, we developed an incremental modeling approach — a methodological strategy common in mathematics and physics but still rare in geroscience that progressively incorporates biological aging factors upon a baseline non-aging model. We employed reliability theory^56^, which conceptualizes organs as critical system components with finite failure tolerance, and conducted extensive literature review to estimate key biological parameters with rigorous validation against empirical bounds.

Thus, this work introduces a framework that explicitly links mechanistic drivers of aging to population-level manifestations, with potential to rank canonical hallmarks by their relative impact through shifts in survival curves. Importantly, our estimates assume: (1) no organ/tissue transplantation; (2) no interventions reducing mutation accumulation; and (3) baseline all-cause mortality fixed at contemporary 30-year-old levels.

Our results suggest that even if all hallmarks of aging were eliminated except for somatic mutations, median human lifespan would reach only 134-170 years — roughly twice the current 76 years. This approximate 2-fold gap in median survival indicates that other hallmarks of aging^10^ likely contribute at least as much to limiting human lifespan as somatic mutations do. Maximum lifespan would range from 210 to 730 years depending on inter-organ dependency structure (627 years under independence). While these estimates substantially exceed current longevity, they fall well short of the optimistic projections of achieving “longevity escape velocity”^75^.

A key insight is that post-mitotic cell types (neurons, cardiomyocytes) act as critical longevity bottlenecks under somatic mutation pressure, with median failure times of 730 years (brain) and 1,236 years (heart) — far shorter than the 14,134-year non-aging baseline. This reflects inexorable mutation accumulation in irreplaceable cells, driving median lifespan from 430 to 169 years for brain aging alone. In stark contrast, proliferating tissues show remarkable resilience. For liver with stem cell support (Model IIIB), no simulated trajectories reached failure within 100,000 years. Even without stem cells (Model IIIA), hepatocytes maintained functionality beyond 37,000 years, as dead cells are continuously replaced by undamaged progeny. The exception is airway basal cells (Model IIIC), which fail at ~ 4,362 years median time due to proliferative exhaustion, suggesting that even replication-competent tissues may still constrain extreme longevity.

Interestingly, applying Model II to both hepatocytes and airway basal cells yields median lifespans approximately equal to BG mortality even without any replication (Extended Data Fig. 3).

This asymmetry between tissue types supports the notion that aging is not uniform across the organism and provides rational basis for prioritizing therapeutic interventions. Somatic mutations alone are unlikely to bottleneck longevity of proliferating tissues like liver and respiratory epithelium, as regenerative capacity neutralizes their impact (provided protective mechanisms like immune surveillance remain fully functional). Conversely, post-mitotic organs such as brain and heart emerge as critical lifespan bottlenecks. Especially, neuronal irreplaceability imposes severe constraints even if other organs are rejuvenated or replaced. In this context, translational cell therapy aimed at neuronal replacement^46^ may represent our only viable path to 150+ year lifespans, even assuming all other hallmarks are mitigated.

Among the limitations, our model cannot be falsified in the conventional sense, as no organism ages exclusively through somatic mutations. However, it permits a critical *sanity check*: predicted lifespans substantially exceed the observed ones (median: 76 years; maximum: 122 years) by approximately 2-fold for median and 2-6-fold for maximum lifespan. This outcome validates our modeling approach: had the predictions fallen below observed values, it would indicate either fundamental flaws in model structure or systematic biases in parameter estimation (e.g., overestimated mutation accumulation rates). Instead, the model suggests that somatic mutations in the four examined organs markedly increase mortality from the non-aging levels, but still cannot fully explain observed human mortality.

Our framework incorporates only four organs and assumes independence — a simplification of real physiological inter-dependencies. The Fréchet bounds (median: 134-170 years; maximum: 210-730 years) provide mathematical limits on how inter-organ dependencies could shift our predictions, but the true dependency structure remains unknown.

Additionally, our model considers only lethal mutations that cause immediate cell death, neglecting potential pathways through which mutations affect tissue function. First, sublethal mutations may impair cellular function without triggering death, gradually degrading tissue maintenance capacity. Second, we assume malignant transformation is effectively cleared by immune surveillance, thereby excluding cancer as a failure mode. Third, and perhaps most importantly for proliferating tissues, we do not model clonal expansion — the preferential proliferation of mutant cell lineages that outcompete other cells. Clonal hematopoiesis, for instance, is prevalent in aging blood and immune tissues and is associated with increased mortality risk, but similar expansion in solid tissues like bronchi and liver is spatially constrained to a small area (e.g., liver nodule)^76^. Therefore, its impact on organismal health is less pronounced, but may still be significant when considering chronic inflammation and cancers.

Accordingly, our outputs should be interpreted as *estimates* grounded in experimentally measured mutation rates, empirically derived cell division kinetics, functionality thresholds, and parsimonious assumptions regarding failure dynamics — analogous to how sample means estimate population parameters.

In perspective, this work establishes an extensible, quantitative foundation for modeling how individual aging mechanisms constrain lifespan. By demonstrating that somatic mutations alone would permit median lifespans of 134-170 years — substantially above current values yet well below the non-aging baseline — we provide an empirical reference against which other hallmarks can be compared. Ongoing efforts profiling somatic mutations across tissues^77^ and species will enable incorporating additional organs and cross-species validation, which may reveal further lifespan-limiting bottlenecks. Parameter estimation would significantly benefit from *in vivo* experiments on replication rates, mutation lethality, and proliferation limits.

Our current Model III incorporates the Hayflick limit for cell proliferation, which can be viewed as a proxy for telomere attrition, another aging hallmark. However, telomere shortening in somatic cells is not aging *per se* — it prevents malignancy by inducing senescence, after which cells should be cleared by immune surveillance^78–80^. Age-related immune decline causes senescent cell accumulation^78,80^, concurrent with stem cell telomere attrition contributing to niche exhaustion^78,81^, though exact pathways remain unclear. Telomere attrition can be accelerated by other hallmarks like mitochondrial dysfunction^82,83^, highlighting the potential interplay for future modeling.

Integrating all hallmarks (mitochondrial dysfunction, proteostasis loss, epigenetic alterations, etc.) requires substantially greater complexity, encompassing extracellular matrix and intercellular networks^84^. However, our work provides a critical first step towards dissecting aging into quantifiable mechanistic components. Ultimately, incorporating other major aging factors could pave the way to a comprehensive, mechanistic theory of aging. Achieving it would require coordinated efforts across multiple research groups — a challenging but increasingly attainable prospect in the near future.

## Methods

### Model I: Constant hazard rate

Survival function *S*(*t*) is defined as the probability that an individual survives beyond the age of *t* years — in other words, it is the probability to die at age *τ* which is later than *t*, therefore:

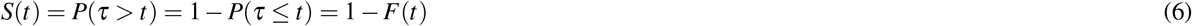

where *F*(*t*) = *P*(*τ* ≤ *t*) is equivalent to the fraction of population that has died by age *t*.

Hazard function *λ* (*t*) is the instantaneous rate of death at age *t*, conditional on having survived to this age *t*. In other words, it is the probability that an individual dies within time interval Δ*t*, given that they survived until *t*, divided by this interval Δ*t*:

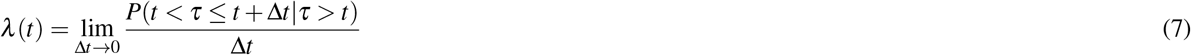

To calculate hazard rates for each age point, we estimated empirical survival probabilities using life-table data obtained from the Human Life-Table Database (HLD; https://www.lifetable.de). Our analysis includes nine geographically diverse populations (United States, Russia, China, Germany, Israel, India, Mexico, South Africa, and Malaysia) across all available years for each country.

Let *t*_1_ *< t*_2_ *<* · · · *< t*_*k*_ be the times of observed death events in a certain country. Empirical survival probability for this country according to the Kaplan-Meier estimator^85^ is:

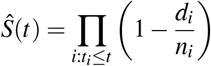

where *d*_*i*_ is the number of deaths at each *t*_*i*_ and *n*_*i*_ is the number of individuals at risk (still alive) just before *t*_*i*_. With the help of a common formula from survival analysis^86^, we can obtain hazard rate as:

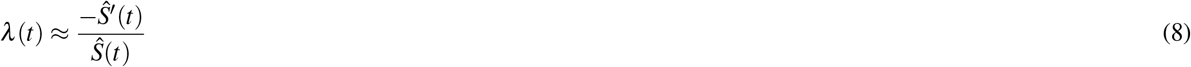

where − *Ŝ′*(*t*) is equivalent to probability density function (PDF) of survival *f* (*t*) = *F*^*′*^(*t*), and is essentially the probability of dying between time points *t* and *t* + Δ*t*. The derivative *Ŝ′*(*t*) was computed from *Ŝ*(*t*) using `numpy.gradient` from the *numpy*^87^ Python package.

If we freeze hazard rate *λ* (*t*) at its value for a certain age *t* and denote it as *λ*, then we can derive the survival function *S*(*t*) for Model I from equations (6), (7), and (8) as:

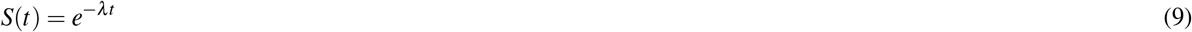

Within this constant mortality framework, we can calculate the maximum and median lifespans *t*_max_ and *t*_median_ as times at which the expected number of survivors from the initial human population of size *N* (assumed to approximately equal 8 billion people^66^) would be reduced to 1 person or to half the initial population, respectively. These times can be found by solving the equations *S*(*t*) = 1*/N* and *S*(*t*) = 1*/*2 for *t*, yielding:

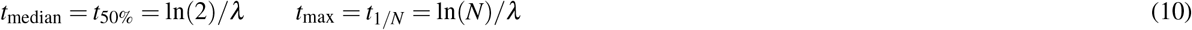

Survival function for the US population was utilized as background (BG) survival function for all further models, hereafter referred to as *S*_BG_(*t*). Hazard rate at age 30 calculated for this population is, accordingly, referred to as background mortality rate *λ*_*BG*_.

### Model II: Cell death

We begin our incremental construction of the aging model by incorporating the phenomenon of cell death driven by somatic mutations. It can be modeled as a simple exponential decay process that acts independently on each cell of a given organ. When cell population falls below a critical threshold, the entire organ and, therefore, organism dies. This scenario can be captured by combining the baseline mortality introduced in Model I with an additional, somatic mutation-specific mortality component, which we introduce below. The model is based on the following assumptions:

- Cell death occurs only due to lethal somatic mutations.
- Mutations cannot be repaired after a certain small time period *dt* passes.
- Mutations occur and kill independently of each other.
- Lethal mutations act instantaneously (no delayed or cumulative effect on the same cell).
- There is no regeneration or replacement of lost cells (decay-only model).

We model cell population *X* = *X* (*t*) change over time as:

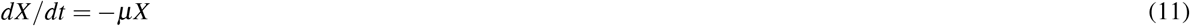

where *µ* = *µ*_0_ · *p*_*lethal*_ is *per-cell hazard rate* equal to the product of *µ*_0_ — the accumulation rate of somatic mutations measured as number of mutations (SNVs and indels) per cell per year, and *p*_*lethal*_ — the probability of a single mutation to be lethal for a cell. The calculation of *µ*_0_ and *p*_*lethal*_ is described in greater detail in sections Estimating accumulation rates *µ*_0_ of somatic mutation burden and Estimating probability of lethal somatic mutations. We set the initial number of the cells to *X* (*t* = 0) = *X*_0_ = *K*, with *K* denoting *total cell capacity* for a given organ.

Individuals are allowed to vary in organ capacity *K* (some people have larger or smaller organs) and the per-cell hazard rate *µ* (some accumulate damage faster or slower), both of which are treated as random variables across the population.

The closed-form solution for this model is

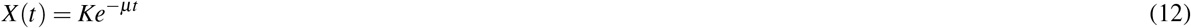

#### Deriving population survival function for Model II

To derive the population-level survival function from individual cell dynamics, we account for inter-individual variation in organ capacity and mutation rates. While individual cells follow exponential decay (*X* (*t*) = *Ke*^−*µt*^), translating this to population mortality requires integrating over heterogeneity in initial conditions, which makes the calculations rather complex.

We start by denoting a critical cell population threshold as *X*_*crit*_. When *X* (*t*) ≤ *X*_*crit*_, the organ can no longer maintain function and the organism dies. Solving *X* (*t* = *t*_*crit*_) = *X*_*crit*_ gives the time-to-organ-failure for an individual with parameters (*K, µ*):

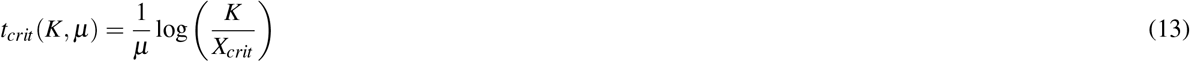

However, instead of asking “when does cell count hit the threshold?”, we can reformulate the problem by asking: “How large must an organ have been at birth to be still alive at age *t*?”

An individual with organ capacity *K* and per-cell mutation hazard *µ* dies when *X* (*t*) = *X*_crit_. Inverting this relationship, we obtain the minimum initial capacity required to survive until *t*:

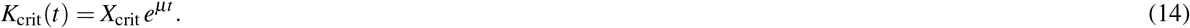

Thus, an individual *i* (with a fixed pair of *K* and *µ*) still survives despite their mutation burden at time *t* if and only if their initial capacity exceeded this threshold:

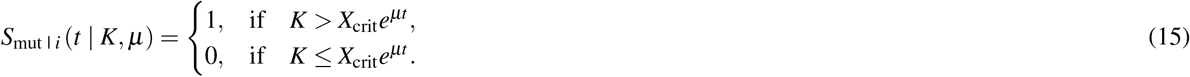

To obtain population survival, we marginalize over the inter-individual distributions (probability density functions, PDFs) of organ capacity *ϕ* (*K*) and lethal mutation rate *ξ* (*µ*). Population survival is therefore an integral over the joint distribution of *K* and *µ* (assuming *K* and *µ* are independent across individuals):

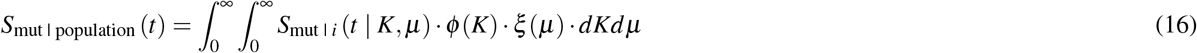

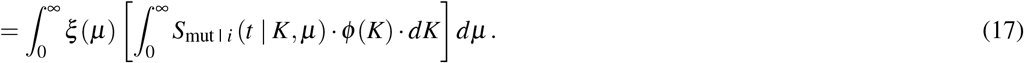

Because death occurs for *K* values below *X*_*crit*_*e*^*µt*^, we can alter the integration limits to include only *K* values above it:

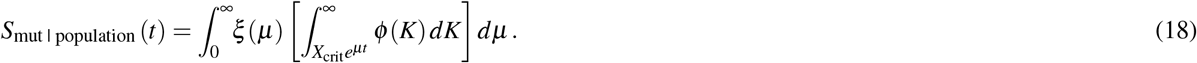

For a fixed *µ*, the inner integral becomes the probability of survival, and it is simply the probability that the initial capacity exceeds the capacity threshold:

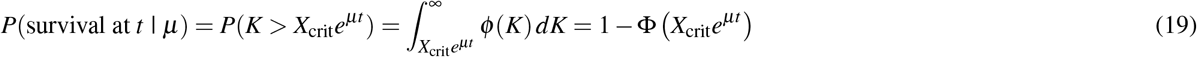

where Φ(…) denotes the cumulative distribution function (CDF) of organ capacity.

To account for the variability in *µ* — some people’s cells die faster, so they need even larger organs to survive to the same age, — we calculate the expectation (i.e., average) of this probability over the distribution of mutation rates by:

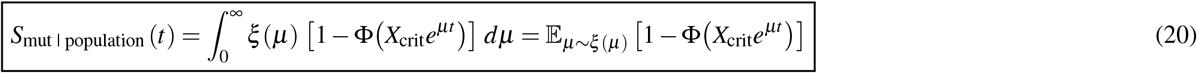

This expression has a natural interpretation: for each mutation hazard *µ*, only individuals with sufficiently large initial organ capacity survive to time *t*; the population survival is the average of this fraction across all *µ* values.

Assuming independence between somatic mutation-driven population mortality and background (BG) hazards, the full survival function is the following:

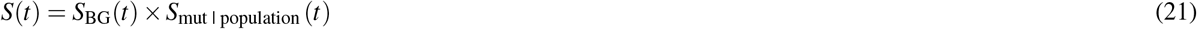

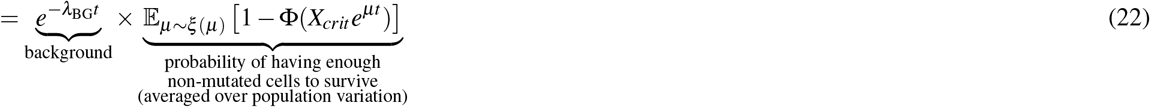

This formulation captures how population survival emerges from the interplay between initial capacity heterogeneity (variation in *K*), aging heterogeneity (variation in *µ*), and the exponential accumulation of cellular damage.

Since analytical solution for *S*(*t*) is intractable, we employed Monte Carlo simulation with 10,000 iterations. For each iteration, we sampled { *K*_*i*_, *µ*_*i*_ } from log-normal prior distributions *ϕ* (*K*) and *ξ* (*µ*) (parameterized further below in sections Estimating accumulation rates *µ*_0_ of somatic mutation burden and Estimating other model parameters for each organ), computed the survival function, and averaged across samples. Maximum and median lifespans were obtained by searching for the first entry where *S*(*t*) ≤ 1*/N* and *S* ≤ 1*/*2, respectively.

### Model IIIA: Cell death and replication of somatic cells

We next add the ability to replicate into our framework in order to model proliferating cell types. To do it, we rely on the following assumptions:

- All assumptions from Model II, but also:
- Cells can replicate, each giving rise to 2 daughters of same cell type. The net gain in cell number from one division is therefore +1. Maximum division rate is *r* (measured in divisions per year).
- Division is limited by available space and resources presented by population capacity (*K*), therefore *replication rate ρ*
- should depend on the current *x* = *X/K* ratio.
- A cell’s ability to divide is proportional to its remaining *replicative (proliferative) potential P* measured in number of divisions left.
- A fresh new cell starts with *P* = *H* total divisions a cell of a given type can undergo (also known as the *Hayflick limit*); cells with *P* = 0 become senescent and can no longer divide, thus making no addition to the current cell count. Over time, potential decreases at rate *ψ*.
- Senescent cells are cleared sufficiently fast by the immune system, without any influence on the neighboring cells (because there is no accumulation of senescent cells and inflammaging in our model).

To create this model, we employ a system of two equations:

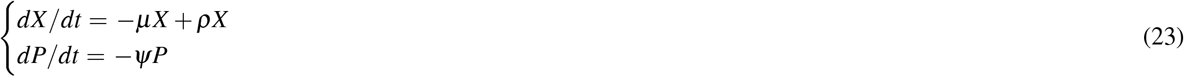

Where

*µX* = number of cells died from somatic mutations,

*ρX* = number of cells produced due to replication, and

*ψP* = loss of proliferative potential due to replication, averaged across population.

#### Modeling replication rate depending on current population and proliferative potential

To model *ρX*, we choose the simplest law of sigmoidal population growth, which models proliferation as a function of current population fraction *x* = *X/K* via a simple *logistic gate function g*(*x*) = (1 − *X/K*) = (1 − *x*). The replication rate *ρ* is proportional to this gate *ρ* ~ *g*(*x*) with a coefficient *r*, that is: *ρ* ~ *r* · *g*(*x*) = *r* (1 − *x*). Thus, when *X* → *K* (cell count approaches organ capacity), then *x* → 1 and *ρ* → 0 (no replication). As *X* falls down close to zero, so that *x* → 0, *ρ* increases up to *r*. Hence, *r* represents *maximum rate* at which the cells of a given type can proliferate (expressed in number of divisions per year). If we consider the *X* (*t*) curve starting from *X*_0_ = 1 (population growth from 1 cell), then its *inflection point* would be where the rate of change of the growth rate switches from speeding up to slowing down (i.e., *X* (*t*) acceleration is zero) — equivalently, where the absolute growth *dX/dt* is largest. Because we asssumed *ρ* ~ *r* (1 − *X/K*), then the inflection point would lie at *x*_infl_ = 0.5 (half of organ capacity).

To introduce proliferative potential, we multiply *r* (1 − *X/K*) by the remaining fraction of replicative potential *P/H* which decreases from 1 to 0, depending on how many cellular divisions have passed. *P/H* = 1 for new cells at *t* = 0, later becoming *P* = 0 and turning proliferating cells into the senescent ones. Therefore, the division rate — in other words, the fraction of cells that divide and increase current population due to replication across *dt* — becomes (Extended Data Fig. 8):

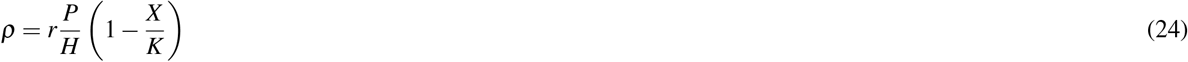

#### Deriving the rate of proliferative potential change

To describe how average proliferative potential changes over time *dP/dt* = −*ψP* (where *ψ* represents the rate of potential loss), we introduce *total potential* Ψ — that is, the number of divisions left across the whole current cell population: 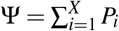. Since *P* is average potential, all cells have the same amount of it. Thus, Ψ = *P* · *X*.

When an *i*-th cell dies, the system loses its *P* remaining divisions, in average: 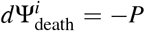. Then, net loss of Ψ due to cell death over *dt* is *d*Ψ_death_*/dt* = −*P* · *µX*. When an *i*-th cell divides, the system “loses” one mother cell with its *P* divisions and “gains” two daughter cells, each with *P* − 1 divisions left. The net change in Ψ per cell division is 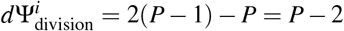. Over *dt*, the number of divided cells is *ρX*. Summing up all divisions that occur over this period, net change of Ψ becomes 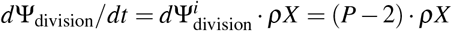. Combining Ψ loss from both death and division, we get

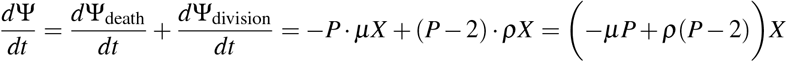

Since Ψ = *P* · *X*, then, according to the product rule for derivatives,

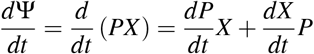

We can rearrange it to solve for *dP/dt*:

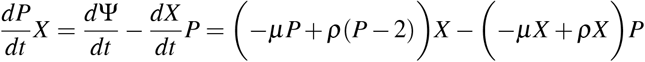

Dividing the whole equation by *X* and expanding the result, we obtain

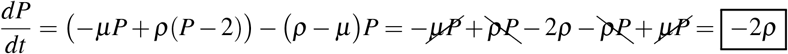

Remembering that *dP/dt* = − *ψP*, we can now derive *ψ* (which is the proportion of potential lost on average across all cells over a given year) as follows:

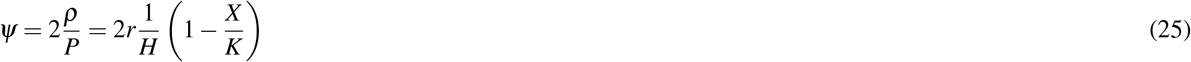

Since *r* = [N divisions/year] and *H* = [N divisions], the *r/H* coefficient becomes [1*/*year]. The (1 − *X/K*) part is dimensionless, therefore *ψP* = [1*/*year], yielding loss of proliferative potential averaged across *dt* = 1 year, as we desired.

#### Full system of equations for Model IIIA

Combining the derviations above, the resulting system for Model IIIA (somatic only) becomes:

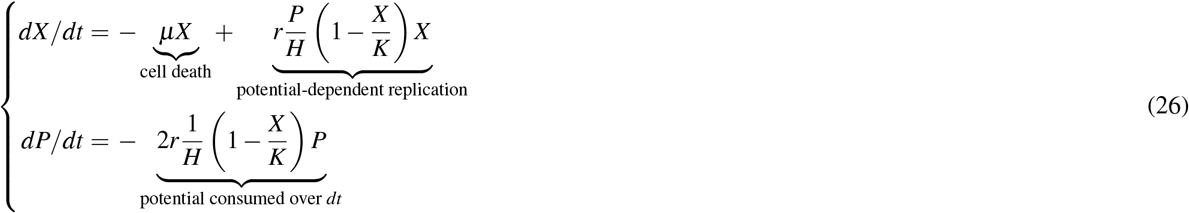

### Model IIIB: Death and replication of somatic and stem cells

After birth, most human tissues harbor a population of progenitor cells which replenish the somatic cell pool, preventing a tissue from the Hayflick-determined lifespan. Historically, these cells have been called various synonyms, depending on the tissue of origin: adult stem cells, somatic stem cells, resident stem cells, progenitor cells, etc. For convenience, in this work we refer to all of them simply as stem cells (SCs), because we do not consider the germline for our modeling. SC proliferation is a continuum^88^: in some tissues, SCs proliferate constantly, in order to support the fast turnover of their somatic descendants, the most prolific examples of which are hematopoietic stem cells producing blood and immune lineages^89^, intestinal crypt cells sustaining intestinal epithelium lining^90^, epidermal stem cells responsible for skin renewal^91^. Other SCs mostly lie quiescent and become activated only after a serious injury occurs, so there is urgent need to replace the dead somatic cells and restore organ function: for example, muscle satellite cells, adipose stem cells, etc.^88^ Some other tissues are virtually devoid of SCs and are, supposedly, not replenished (we employ Model II for those).

Modeling SCs that belong to rapid turnover niches (such as hematopoietic stem cells) appears challenging in our framework, because these are especially prone to clonal expansion, meaning that somatic mutations can hardly be assumed independent, and that an organ can deteriorate functionally, while still being able to sustain its cell counts. Moreover, the plasticity of these SCs allows them to choose their fate dynamically, such that the proportions between self-renewing division and differentiation into various somatic lineages (multipotency) might change dynamically as well, making the models ever more cumbersome. For the quiescent SCs, on the other hand, their data on somatic mutagenesis and proliferation parameters are markedly more scarce. As a middle ground, we chose liver progenitor cells (LPCs, also known as oval cells) which are known to be bipotent and differentiate into either hepatocyte or cholangiocyte populations of adult human liver: cholangiocytes are epithelial cells that line out the bile ducts, while hepatocytes are the main players responsible for blood detoxification, lipid formation, gluconeogenesis, bile secretion, protein synthesis and storage, etc.^92–94^. With so many jobs, hepatocytes are under constant stress, especially considering their key role in toxin removal. However, they can apparently proliferate quite a handful of times, so that the LPCs are normally quiescent throughout human life, unless severe liver damage is done: e.g., the removal of a sizable bulk of liver cells (*>*50% and probably even *>*70% hepatocyte loss) or large amounts of toxins pouring into the bloodstream^92,95,96^.

The magnificent proliferative capacity of hepatocytes has recently been attributed to the heterogeneity within their ranks: in mouse livers, there are hepatocyte subtypes which express high levels of telomerase (*TERT*), thus maintaining their telomere length without senescence^97^. The members of this *TERT*^*high*^ species represent approximately 30% of murine hepatocyte population, reside further away from the ducts, and are likely the ones responsible for restoring the common hepatocyte pool by differentiating into their more abundant, *TERT*^*low*^ siblings which either reached their replicative senescence or succumbed to external damage. In adult humans, however, *TERT* expression has not been reported^98,99^. Other putatively interesting hepatocyte subtypes have been described in humans and mice^93,100–102^, but incorporating all of them into our model is currently unfeasible, because there is no data on replication rates and mutagenesis with such subtype-level resolution. Taking all that into account, we treat hepatocytes as a single population, relying on the average estimates of their parameters.

Our assumptions for this model are:

- As in Model IIIA, but also:
- There is a *population of stem cells Y*; maximum *stem cell capacity* per organ is *Q*.
- Stem cells die over time due to somatic mutations, as well as somatic cells, albeit at their own rate *µ*_*y*_.
- Stem cells can undergo three types of division: either *symmetrical self-renewal*, which gives rise to 2 daughter stem cells, or *symmetrical differentiation*, producing 2 daughter somatic cells, or *asymmetrical division*, which yields 1 stem and 1 somatic daughter. Thus, stem cell division adds new cells to both somatic (*X*) and stem (*Y*) populations.
- In sum, the fractions of the respective division types compound to: *f*_*yy*_ + *f*_*xx*_ + *f*_*xy*_ = 1.
- *Stem cell division rate ρ*_*y*_ depends both on *X/K* and on *Y/Q*.
- Proliferative potential of stem cells is constantly full (that is, stem cell death and niche exhaustion can occur only due to somatic mutagenesis, all other mechanisms of aging do not manifest).
- Stem-derived somatic cells bear full proliferative potential, replenishing that of the current somatic population.

To incorporate SCs into our modeling, we need to add a few terms and a separate equation:

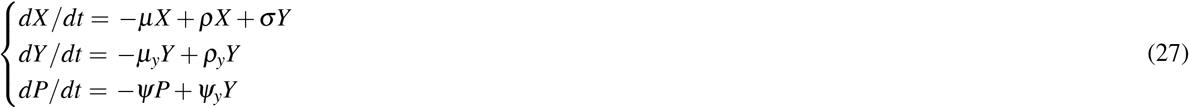

where *σY* = number of new somatic cells (with *P* = *H*) derived from stem cells,

*µ*_*y*_*Y* = number of stem cells died due to mutations,

*ρ*_*y*_*Y* = number of stem cells produced due to their division, and

*ψ*_*y*_*Y* = proliferative potential of somatic cells replenished by stem cell differentiation.

#### Modeling rates related to stem cell death and division

As for somatic cells, *µ*_*y*_*Y* is calculated using baseline mutation accumulation rate *µ*_0 | *y*_ and the probability that a given mutation becomes lethal for a stem cell:

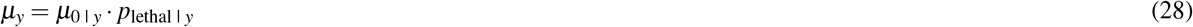

To model both *σ* and *ρ*_*y*_, we can employ a formula similar to somatic replication: *r*_*y*_ · *g*(*x*), where *r*_*y*_ is maximum stem cell replication rate, and a sigmoid gate *g*(*x*) is responsible for the non-linear population growth between homeostasis and recovery. Given that LPCs engage in replication only after a significant portion of somatic cells is gone (*x* = *X/K* is low), we can employ a more flexible, *Richards-type logistic gate* (in one of its reparameterizations)^103^: *g*(*x*) = (1 − *x*)^*m*^. To find the inflection point (the point of zero acceleration), we can solve the *d*^2^*X/dt*^2^ = 0 equation and obtain:

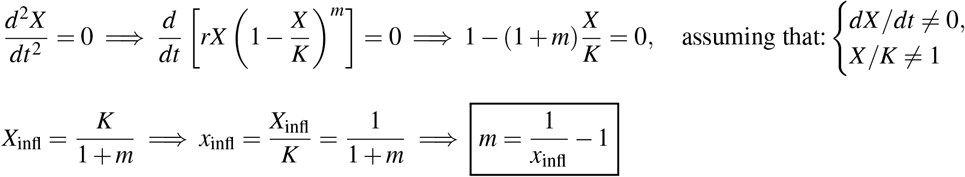

Thus, if we want to set the inflection point to *x*_infl_ = 1*/*3 (maximum population growth at 1*/*3 of organ capacity), then we can use 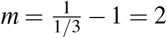. Evidently, when *m* = 1, the Richards-type logistic becomes the simple logistic *g*(*x*) = 1 − *x*, with *x*_infl_ = 1*/*2 (maximum population growth at half of organ capacity), which we use for all other cell types except for LPCs.

However, LPCs proliferate not only after large hepatocyte loss (when *x* = *X/K* is low), but also in response to toxin-induced damage to the LPC pool (when *y* = *Y/Q* is low)^92,96^, so the proliferation law for LPCs has to reflect both hepatocyte density and LPC niche occupancy. We can modify our Richards-type gate as a function *g*(*x, s*) of both *x* and *s* in various ways, the simplest of which are multiplication (logical AND-like) and addition (logical OR-like):

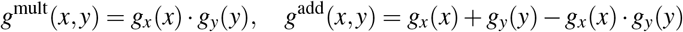

Because the addition lets either hepatocyte loss OR low stem occupancy trigger high proliferation (Extended Data Fig. 10, we chose the additive gate to represent LPC proliferation in our model:

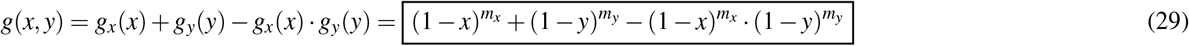

where *m*_*x*_ and *m*_*s*_ are shape parameters that describe how sharply each gate falls off as its corresponding population fraction (*x* = *X/K* or *y* = *Y/Q*) increases. We will further refer to *g*(*x, y*) as *g* for simplicity.

The resulting product of *r*_*y*_ · *g* is *total stem division rate* 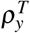. To find which proportion of this rate yields actual SCs (*ρ*_*y*_) and not the differentiated ones (*σ*), we should adjust total rate 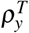 by the appropriate *balance* (denoted as *γ*) between the mentioned division types an SC can undergo. Whichever is the fraction of asymmetric divisions, the stem pool neither loses nor gains any cells from it (0 · *f*_*xy*_). The number of SCs produced (or lost) due to their division is therefore a balance between the fraction of self-renewing divisions (+1 · *f*_*yy*_) and the fraction of differentiation events (−1 · *f*_*xx*_), yielding:

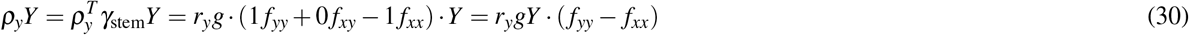

Somatic cells can arise either as 2 daughters of a symmetric differentiation (+2 · *f*_*xx*_), or as 1 daughter of an asymmetric one (+1 · *f*_*xy*_), while the effect of symmetric self-renewal on somatic population is zero (0 · *f*_*yy*_). Therefore, net somatic gain is *γ*_soma_ = (2 *f*_*xx*_ + 1 *f*_*xy*_ + 0 *f*_*yy*_), and

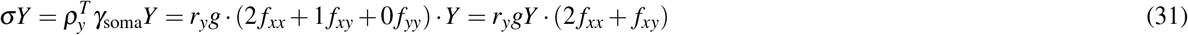

#### Deriving the rates of proliferative potential change considering stem cells

Finally, to obtain a formula for proliferative potential replenished by the fresh, stem-derived somatic cells, we once again resort to the concept of total potential Ψ = *P* · *X* (number of divisions left across the whole population). For somatic population only, Ψ change per division was either due to cell death (by 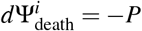), or due to cell division 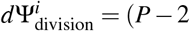). This time, we have the third player — stem-driven replenishment. When an *i*-th cell differentiates from an SC, it increases total potential by 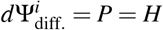. Over *dt*, the number of newly differentiated cells is *σY*. Thus, total potential change due to differentiation over this time is 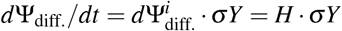. Combining the three equations yields

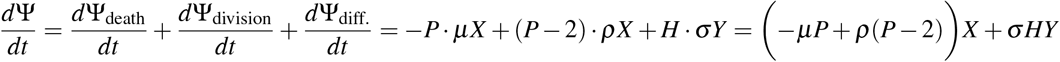

As in the somatic-only model, Ψ = *P* · *X*, allowing us to derive:

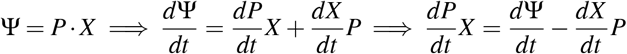

Inserting the combined equation for *d*Ψ*/dt* yields:

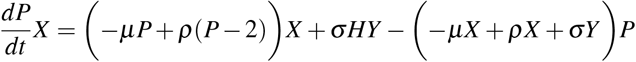

As before, we divide the whole equation by *X*, expand the result, and obtain

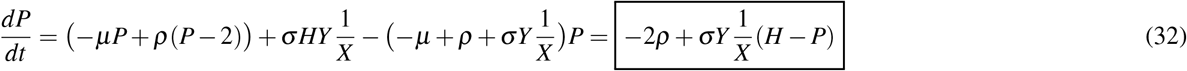

Remembering that *dP/dt* = −*ψP* + *ψ*_*y*_*Y* and that −*ψP* = −2*ρ*, we can now derive *ψ*_*y*_ (the proportion of potential gained from newly differentiated cells per year) by replacing *σ* with its definition and dividing 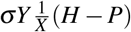 by *Y*:

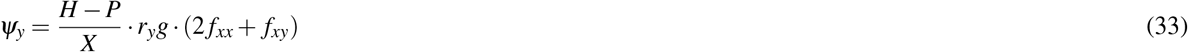

The coefficient 1*/X* can be called a dilution coefficient: it adjusts the input of new cells into the existing potential by dividing it over the current population. Thus, if *X* is high, then the effect of stem-derived cells on average potential is tiny. On the contrary, when *X* drops, every new differentiation markedly impacts the population’s proliferative potential.

Similarly with the (*H* − *P*) multiplier, which can be thought of as the “room for improvement” or as the magnitude of potential deficit — it is the difference between the maximum possible potential (*H*) of a new cell arriving from the stem pool and the current average potential (*P*) of somatic population. When somatic cells are young, *P* is close to *H*, meaning that (*H P*) → 0. The arrival of newly differentiated cells does not make a huge impact on average potential, because current population is already near-perfect. As somatic cells grow old, *P* → 0, meaning that (*H* − *P*) → *H*. The potential deficit is enormous: every new cell with *P* = *H* provides a massive boost to the average potential.

Thus, the (*H* − *P*)*/X* factor creates the situation such that the restorative effect is at its strongest when the tissue is most exhausted. Curiously, this factor arises by itself from our simple principles, recall that we did not assume such dilution or deficit properties for our equations from the start.

#### Full system of equations for Model IIIB

Summarizing equations (28), (29), (30), (31), (32), and (33), the resulting system for Model IIIB (somatic & stem) becomes:

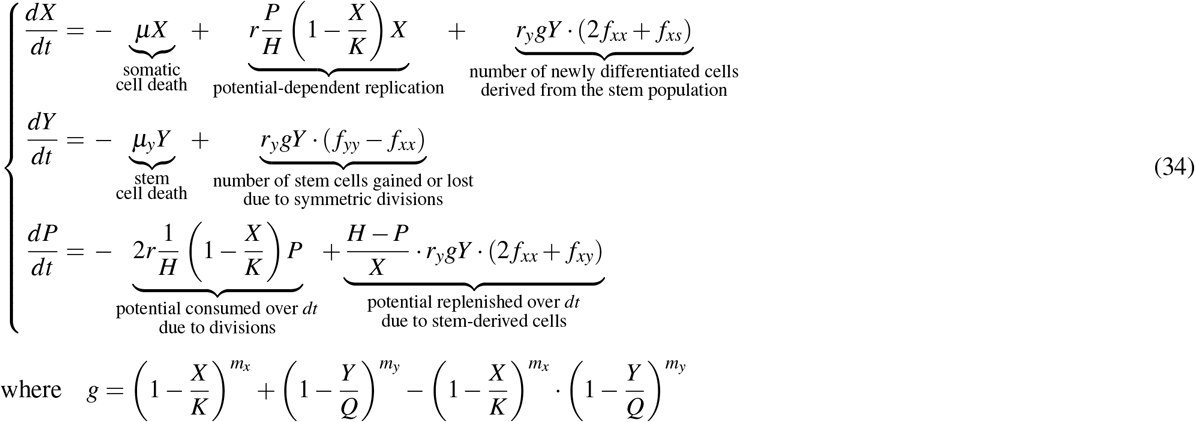

### Model IIIC: Death of differentiating stem cells with limited proliferative potential

Among the cell types whose single-cell profiles of somatic mutagenesis were obtained, there is one peculiar species: of human bronchial basal cells (hereafter referred to as BBCs, also called airway basal cells and HBECs). These multipotent cells give rise to the many cell types of lung airway epithelium, playing the key role in its homeostasis and regeneration, which makes them a valid member of the somatic stem cell family. However, their proliferation was also shown to be finite in culture: earlier studies at atmospheric 21% O_2_, together with a ROCK (Rho-associated coiled-coil kinase) inhibitor resulted in BBC senescence at 80 population doublings^104^, while milder, 2% O_2_ conditions allowed them to survive 200+ doublings^105^. Even though the *in vivo* data is scarce, the fact that BBCs require exceptionally mild conditions to continue proliferating points us towards assuming that their division *in vivo* is also limited.

Recently, it was shown that there are actually two morphologically identical subpopulations of airway basal cells (in mouse trachea), one of which is basal stem cells, while the other is their luminal progenitors^106^. Only the progenitors can differentiate further, thus serving as a transient cell type between the stem cells and the secretory or ciliated cells. However, whether the same two subtypes exist in human airway epithelium and, if yes, then what their respective mutation rates are, remains unresolved. Therefore, we treat airway basal cells as a single population, relying on the average estimates of their parameters (as with hepatocytes).

Our assumptions for this model are:

- As in Model II, but:
- There is a population of BBCs *B*; maximum BBC population per organism is *K*.
- BBCs die only from somatic mutations at rate *µ*_*b*_, as with other cell types in our modeling.
- BBCs can undergo the same types of stem cell division mentioned earlier: either symmetrical (self-renewal or differentiation) or asymmetrical. BBC division can both reduce and produce their population (*B*).
- The fractions of BBC division types compound to: *f*_*bb*_ + *f*_*xx*_ + *f*_*xb*_ = 1.
- Proliferative potential *P* decreases with each BBC division (of any type) at rate *ψ*_*b*_.

To model BBC behavior, we need to re-adjust our system:

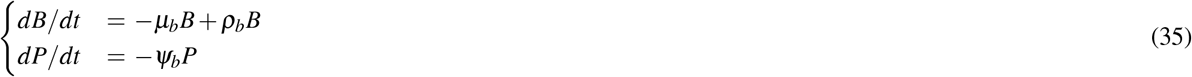

where *µ*_*b*_*B* = number of BBCs died due to mutations,

*ρ*_*b*_*B* = number of BBCs produced due to their division, and

*ψ*_*b*_*P* = proliferative potential of BBCs consumed by their division.

As with other cell types,

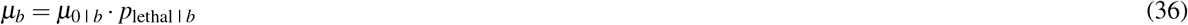

#### Modeling BBC replication rate

The rate of division *ρ*_*b*_ can be modeled with the similar *r*_*b*_ · *g*_*b*_ as for other stem cells, adjusted by the current fraction of divisions left *P/H*. As there is little information on BBC-specific patterns of population growth, we assume a simple sigmoid (1 − *B/K*), where *K* is total BBC capacity of human airways. Combining maximum division rate *r*_*b*_, sigmoid growth based on current population size (1 −*B/K*), and the proliferative potential coefficient *P/H*, we obtain total division rate:

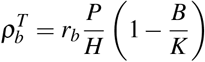

Since only a fraction of this rate yields BBCs, while the rest goes away to differentiation, we need to adjust 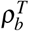 by the fraction of BBC-producing divisions among all divisions, denoted as *γ*_*b*_. As with model 3, the BBC pool neither loses nor gains any cells from asymmetric divisions (0 · *f*_*xb*_), and the net gain of BBCs due to their division is a balance between self-renewal (+1 · *f*_*ss*_) and differentiation (−1 · *f*_*xx*_), yielding:

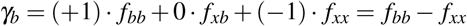

Combining total division rate 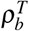 and this fraction of BBC-producing divisions *γ*_*b*_, we obtain BBC-producing division rate:

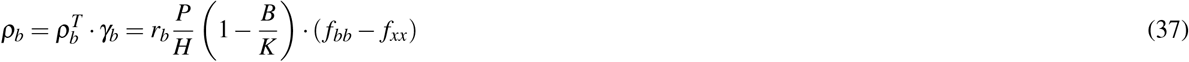

As for *dP/dt*, we resort once again to the concept of total system potential. The BBC system has no external source of new potential — it is only consumed, even though BBCs are multipotent (stem-like). The degradation of average potential *P* should be proportional to the total rate of division events occurring in the population, as each division consumes BBC potential.

#### Deriving the rate of proliferative potential change in BBCs

Due to cell death, total potential Ψ changes over time by *d*Ψ_death_*/dt* = −*P*· *µ*_*b*_*B*. The change of total potential due to divisions can be found through multiplying the net change of potential per division 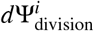 by the total number of divisions over time 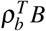. Due to any division, the BBC pool loses 1 mother cell with potential *P* and gains daughter cells, each with potential *P*− 1. If both daughters are BBCs, then 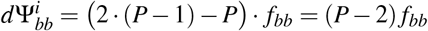, if only one daughter is a BBC, then the amount of potential added to (or, in other words, remained within) the BBC pool is 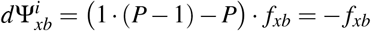. Sym metric differentiation only consumes mother potential, without adding any daughters to the total pool 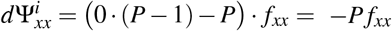. Total potential change due to different division types is

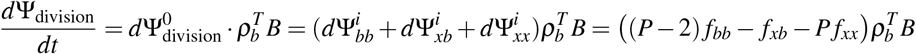

Thus, combining BBC death and division,

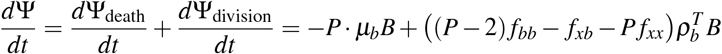

Similarly to the previous models, Ψ = *P* · *B*, allowing us to derive:

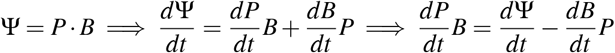

Inserting the combined equation for *d*Ψ*/dt* yields:

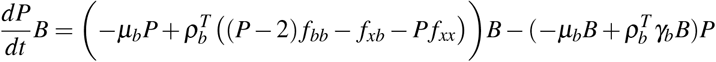

As before, we divide the whole equation by *B*, expand the result, expand *γ*_*b*_, and obtain

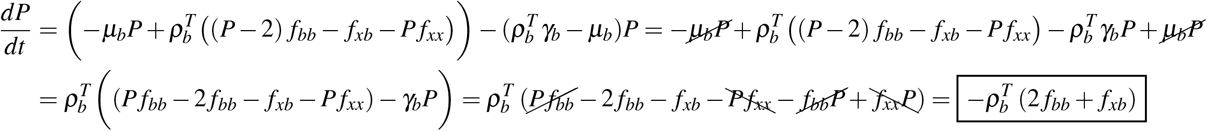

Remembering that *dP/dt* = −*ψ*_*b*_*P*, we can now derive *ψ*_*b*_ (rate of potential loss per year):

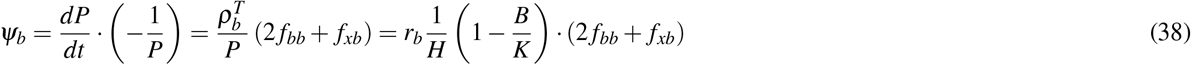

#### Full system of equations for Model IIIC

Summarizing the equations for *dB/dt* and *dP/dt*, the resulting system for Model IV becomes:

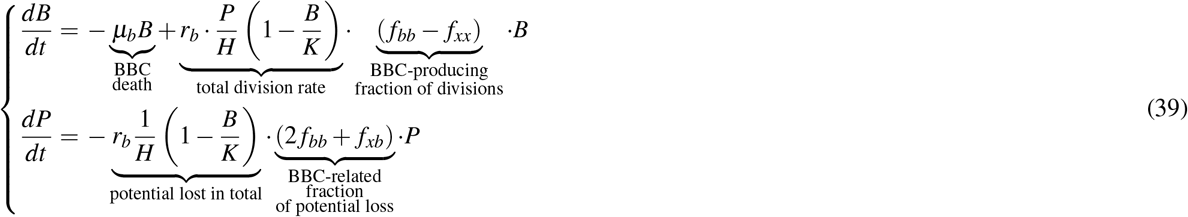

For clearer understanding of the last equation for *dP/dt*, we can expand the second bracket and replace 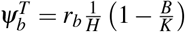:

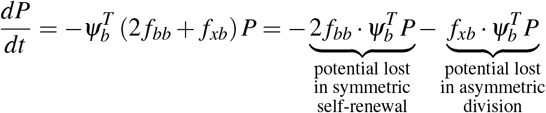

There is no *f*_*xx*_-containing term, because differentiated cells make no change to the average potential of BBCs.

### Processing data on somatic mutations

The landscape of somatic mutations is challenging to profile, as they occur in each cell independently and therefore can hardly be detected in bulk tissue samples. Traditional genomic techniques were able to detect only large-scale chromosomal alterations (e.g., aneuploidy, megabase copy number variations), while the estimation of single nucleotide variants (SNVs) and insertions/deletions (indels) remained unfeasible^107,108^. To overcome these limitations, techniques like single-cell cloning and laser capture microdissection (LCM) were developed. However, their applicability is constrained by tissue type — particularly for non-proliferating tissues. The emergence of single-cell whole-genome sequencing (scWGS) now provides the most robust method for measuring the somatic mutation burden within individual cells, including rare, non-clonal variants^77^. ScWGS almost always requires whole-genome amplification to produce enough DNA for sequencing, which can be done in several ways. Among these, the most widely used are MDA^109^ (multiple displacement amplification) and PTA^110^ (primary template-directed amplification). Although MDA was among the first methods to produce large DNA fragments that yielded good coverage with a relatively straightforward protocol, it suffers from allelic dropout (one allele may fail to amplify), biased genome coverage, and amplification artifacts (artificial SNVs/indels), especially due to late amplification errors and single-strand dropout^110,111^. PTA allows more uniform, accurate, sensitive, and reproducible profiling of somatic mutations than any other existing scWGS approach^70,110,111^. We therefore prioritized PTA-based scWGS datasets for our analysis. For tissues where PTA data were unavailable, we utilized MDA-based datasets as an alternative.

For Model II (only death, no replication), we focused on brain neurons and heart cardiomyocytes. Models IIIA (somatic only) and IIIB (somatic & stem) were employed for liver hepatocytes and liver progenitor cells (LPCs). Model IIIC was run on human bronchial basal cells (BBCs) retrieved from proximal lung airways. Model II was additionally run on hepatocytes and BBCs to check their survival dynamics in the absence of any replication (Extended Data Fig. 3).

For neurons, we obtained Variant Call Format (VCF) files, metadata, and/or other data tables from the studies by Luquette et al.^70^, Ganz et al.^71^, and Motyer et al.^72^. From these, we selected only neuronal samples sequenced with PTA. For heart, we downloaded supplementary data from the PTA-based pre-print study on left ventricle cardiomyocytes by Choudhury et al.^73^. Data on liver hepatocytes and LPCs came from the study by Brazhnik et al.^112^, while BBCs were profiled by Huang et al.^113^. For both liver and lung, we retrieved the respective VCFs (genome build hg38) from SomaMutDB — a database of somatic mutations^114^, while sample metadata were downloaded from the supplementary sections of these articles. From all datasets, we selected samples exclusively from healthy controls. For lung tissue, we further restricted to non-smokers. We excluded outliers with exceptionally high or low mutational burden, as well as samples with extended post-mortem intervals (PMI ≥ 50 hours). The resulting outlier sample list is: “`4638-Neuron-4`”, “`CT2_PL_Neu2`”, and “`CT2_PL_Neu3`” (Brain); “`1039_A1`”, “`1039_A5`”, and “`5828_E6`” (Heart); “`N1274_2`”, “`N1274_8A`”, “`N1276_4`”, “`N1407_2`”, “`N1410_1`”, “`N1410_6`”, and “`N1415_3`” (Lung).

A summary of all analyzed somatic mutation datasets and the resulting sample counts is provided in Table 1.

**Table 1.** Datasets of somatic mutations included in the analysis.

| Tissue | Cell type | Source paper | Amplification method | Genome build | Number of samples |
| --- | --- | --- | --- | --- | --- |
| Brain | Neurons | Luquette et al., 2022<sup>70</sup> | PTA | hg19 | 51 |
| Brain | Neurons | Ganz et al., 2024<sup>71</sup> | PTA | hg19 | 4 |
| Brain | Neurons | Motyer et al., 2025<sup>72</sup> | PTA | hg19 | 10 |
| Heart | Cardiomyocytes | Choudhury et al., 2025<sup>73</sup> | PTA | hg19 | 47 |
| Liver | Hepatocytes | Brazhnik et al., 2020<sup>112</sup> | MDA | hg38 | 51 |
| Liver | liver progenitor cells | Brazhnik et al., 2020<sup>112</sup> | MDA | hg38 | 13 |
| Lung | Bronchial basal cells | Huang et al., 2022<sup>113</sup> | MDA | hg38 | 53 |

The hg19-based datasets were converted into the hg38 assembly using Python *liftover* (v1.3.0) package (https://github.com/jeremymcrae/liftover).

### Estimating accumulation rates *µ* _0_ of somatic mutation burden

For datasets from Luquette et al.^70^ and Ganz et al.^71^, we calculated mutation burden per cell from raw VCF files by counting mutation calls and dividing by the reported sensitivity (48.7% and 46.2% for SNV and indel calling in Luquette et al.; 48% and 41% for SNV and indel calling in Ganz et al.). For the remaining datasets, we obtained mutation burden values directly from supplementary materials, which were already sensitivity-adjusted.

Accumulation rates per year of mutation burden per cell were estimated with a mixed-effect linear model accounting for donor-specific affects. Mixed-effects model fitting was performed for each organ separately, and for SNVs and indels separately, using the *statsmodels*^115^ Python package and its function `mixedlm(mut_burden` ~ `age + (1 | donor_id))`, with REML (restricted maximum likelihood)-based estimation turned on. Accumulation rates were retrieved as slope coefficients returned by this function. Slope confidence intervals (CIs) were calculated from regression values assuming normal distribution and using the *scipy*^116^ Python package function `norm.ppf`, at the 95% confidence level.

Unfortunately, indel calls were unavailable from Choudhury et al.^73^ and Motyer et al.^72^. Additionally, Luquette et al.^70^, Ganz et al.^71^, and Brazhnik et al.^112^ did not report per-cell indel burdens, though their datasets (retrieved from supplementary files or SomaMutDB) contained raw indel calls. To estimate indel accumulation rates for brain and liver cells, we scaled raw indel burdens by the ratio of adjusted-to-raw SNV burdens, assuming similar detection sensitivities for SNVs and indels within each sample, then applied mixed-effects modeling. Since cardiomyocytes share post-mitotic, long-lived characteristics with neurons, we hypothesized similar SNV-to-indel accumulation ratios. In the absence of direct cardiomyocyte indel measurements, we inferred these rates by dividing cardiomyocyte SNV accumulation parameters by the neuronal SNV-to-indel ratio.

SNV burdens for LPCs did not show an upward trend with age (slope = − 16.176, 95% CI = [− 83.627, 51.275], P-value = 0.64, intercept = 1021.476), likely due to the limited sample size (10 samples from 3 donors, all below 18 years). In the original article, Brazhnik et al.^112^ explored LPC-derived organoids to compare their SNV accumulation rates with hepatocytes.

Rather than employing the LPC-derived organoid data from Brazhnik et al.^112^, we maintained an *in vivo* framework by finding the hepatocyte-to-LPC ratio of total SNVs accumulated per cell in young (≤ 36 years) individuals (similarly to the analysis by Brazhnik et al.^112^ demonstrated in the paper’s Figure 1C), obtaining values of 1.565 (SNVs) and 1.671 (indels). LPC mutation accumulation rates were thus estimated by dividing the hepatocyte-modeled coefficients by these ratios.

Results of the accumulation rates estimation are demonstrated in Table 2.

**Table 2.** Mutation accumulation rates per tissue according to the mixed-effects modeling.

| Tissue | Mutation type | Slope | CI low | CI high | Slope P-value | Intercept |
| --- | --- | --- | --- | --- | --- | --- |
| Brain | SNVs | 17.489 | 16.112 | 18.865 | 6.0e-137 | 73.699 |
| Brain | Indels | 6.926 | 5.484 | 8.367 | 4.6e-21 | -41.083 |
| Heart | SNVs | 36.369 | 19.519 | 53.218 | 2.3e-05 | 97.973 |
| Heart | Indels | 14.403 | 6.644 | 23.604 | — (no fitting) | -54.613 |
| Liver | SNVs | 52.783 | 36.647 | 68.920 | 1.4e-10 | 572.414 |
| Liver | Indels | 1.158 | 0.721 | 1.595 | 2.1e-07 | 9.209 |
| Liver (LPCs) | SNVs | 33.726 | 23.415 | 44.036 | — (no fitting) | 365.740 |
| Liver (LPCs) | Indels | 0.693 | 0.432 | 0.955 | — (no fitting) | 5.512 |
| Lung | SNVs | 28.519 | 20.725 | 36.312 | 7.4e-13 | 648.586 |
| Lung | Indels | 2.573 | 1.182 | 3.963 | 2.9e-04 | 58.516 |

### Estimating the probability of lethal somatic mutations

To find the lethal mutations rates (*µ*), we adjusted mutation accumulation rates obtained in the previous section (*µ*_0_) by the probability of a mutation to be lethal for a cell of a given type (*p*_lethal_). Estimating this probability is no trivial task, because, to our knowledge, there are no data which could yield these probabilities directly. Therefore, before proceeding with the estimations, we first evaluated the upper and lower bounds as sanity check — any estimated probability should be within this range.

#### Establishing lower/upper bounds for lethal mutation probability

We estimated the lower bound as the probability that a mutation occurs in the coding sequence (CDS) of a single gene whose disruption is lethal for any human cell type. The largest subunit of human RNA polymerase II (*POLR2A* gene) can serve as a representative example of such gene. The CDS of *POLR2A* (the principal transcript: RefSeq NM_000937.5) is 5,913 bp long, encoding a protein of 1,970 amino acids.

For each 3 base long codon, there are 9 possible SNVs. Among 549 total SNVs (61 codons × 9 mutations), 23 are nonsense changes that generate a premature stop codon. Fraction of nonsense SNVs is thus *f*_nonsense_ = 23*/*(61· 9) = 0.0419 = 4.19%.

Given that *POLR2A* comprises 29 exons (as per RefSeq GRCh38.p14 assembly), it also has 28 intervening introns. Every intron is flanked by splice sites: the canonical GT at the 5’ end (donor) and AG at the 3’ end (acceptor). An SNV hitting either of those bases typically abolishes proper splicing of the neighboring exon. Then, the fraction of all SNVs in the coding region that hit essential splice sites is *f*_splice_ = (28 introns · 4 bases per intron)*/*5913 ≈ 0.0189 = 1.89%.

Missense changes make up the largest share of possible SNVs (392 out of 549, which is 0.714 = 71.4%), but only a subset of them will actually knock *POLR2A* out of function. Deep mutational scanning (DMS) — where every possible single-amino-acid substitution is assayed in pooled functional screens — and other experiments find that on the order of 20-30% of missense variants measurably impair protein activity in enzymes and other essential domains^117,118^.

The human genome comprises ~ 3.1 · 10^9^ base pairs (as per RefSeq GRCh38.p14 assembly), yielding ~ 6.2· 10^9^ base pairs for a diploid cell. Combining the fractions of nonsense (stop-gain) mutations, essential splice-site mutations, and deleterious missense variants (taking 20% as the lower bound), we can estimate the lower bound for probability of lethal mutations as follows:

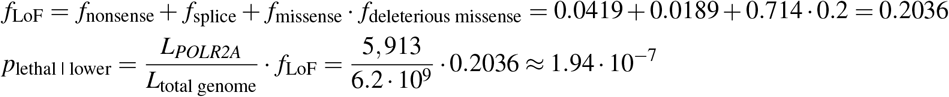

To set the upper bound, we resorted to the SNV burdens obtained in the section on Estimating accumulation rates *µ*_0_ of somatic mutation burden. By age 50, a cell of any analyzed type acquires more than 1000 SNVs (Extended Data Fig. 2). If we assume that, above these, at least 1 is lethal, then there should be approximately 1*/*(1000 + 1) ≈ 0.001 = 0.1% of cells dying each year on average. Summing up across 100 years, we should then observe that ~ 10% of cells per organ are dead by the age of 100 years (purely due to SNVs), which, for instance, contradicts the latest measurements of human cerebral cortex neurons stating that there is no observable neuron loss in cognitively healthy subjects between 25 and 87 years^119^. Therefore, we can safely assume the upper bound for the probability of lethal mutations:

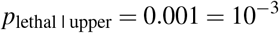

The range between 1.94 · 10^−7^ and 10^−3^ is admittedly wide, but it provides us with the prior boundaries necessary to perform the sanity check of our estimations.

#### Essential genes annotation

Moving on from the single *POLR2A* gene, we next sought other genes that could be essential for cell survival in each analyzed tissue. The concept of gene essentiality is a rather challenging topic, with many caveats and unsolved issues^120–124^. Large-scale studies of genes essential for survival, development, or regeneration are typically performed using CRISPR-based inhibition/activation screens. The *in vivo* human screens of gene essentiality remain ethically unfeasible. Recently, such screens were performed for neurons which had been differentiated from induced pluripotent stem cells (iPSCs)^125,126^. However, the iPSC-derived neurons may not fully replicate gene expression patterns and chromatin features of mature human neurons^127^. Therefore, we relied on *in vivo* mouse screens to determine our panels of essential genes for each cell type, admitting that human essential genes might not be fully captured in the knockout (KO) mouse models^120^.

Neuron-specific^128,129^, cardiomyocyte-specific^130,131^, and liver-specific^132,133^ essential genes were obtained from the respective papers of CRISPR-based screening. Genes passing the p-value threshold of 0.1 after the Benjamini-Hochberg multiple testing correction (as provided in the original papers) were selected. Mouse gene symbols were converted into their human homologs using the *gseapy*^134^ Python package. Since healthy lung-specific essentiality screens remain unavailable to date, we defined lung-essential genes as those reported as essential in > 50% of human lung-derived cell lines in the OGEEv3 database^135^. Each tissue-specific signature of essential genes was united with core-essential human genes (genes reported to be essential across 80+% of all human cell lines) from the OGEEv3 database^135^. The UpSet plot of the resulting signatures and their overlaps is created using the UpSetPlot^136^ Python package and displayed in Extended Data Fig. 6.

#### Annotation by deleteriousness

Mutations do not occur randomly across the genome, and every cell type accrues them in a different way; therefore, we could not assume that the mutations accumulated at rate *µ*_0_ fall in essential and non-essential genomic regions with equal probabilities. Instead, we relied on the empirically observed mutation distributions across the genome.

The probability of a mutation to be lethal for each *j*-th mutation class was modeled as:

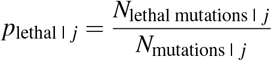

We incorporate two mutation classes in our model: SNVs and indels. *N*_SNVs_ and *N*_indels_ denote the total numbers of SNVs and indels observed per tissue in the same datasets that we used to estimate accumulation rates *µ*_0_ of somatic mutation burden. We estimated lethal mutation counts (*N*_lethal SNVs_ and *N*_lethal indels_) by computing lethality scores *c*_*i*_ for each mutation and summing them:

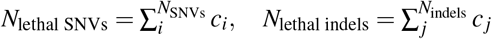

The lethality score *c* is a compound metric which consists of three key factors: 1) whether a mutation falls into a region associated with an essential gene, 2) how deleterious this mutation is, and 3) the probability of this mutation to be haploinsufficient:

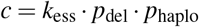

The essentiality coefficient *k*_ess_ takes values 1 or 0 (essential-related or unrelated) and is inferred by intersecting the VEP- and FAVOR-merged annotation (including genes, their upstream/downstream regions, and CAGE promoters/enhancers) of a given mutation with the tissue-specific list of essential genes (see section on Essential genes annotation).

Estimating the effect of a given mutation on its associated gene is another challenging issue, especially for the non-coding regions. To do that, we annotated each mutation using the Ensembl Variant Effect Predictor (VEP)^137^ and the Functional Annotation of Variants – Online Resource (FAVOR)^138^ databases. Then, we calculated *p*_del_ in two steps: first, we aggregated scores from several deleteriousness predictors available in these databases by taking their median. Second, since raw predictor scores do not directly represent the probability of deleteriousness, we applied score calibration.

Pejaver et al.^139^ show posterior probability curves of multiple pathogenicity predictors, from which we inferred that the predictors with the strongest evidence for pathogenicity can be calibrated by a power law *p*_del_ ≈ 0.9 · (score)^6^, with many weaker predictors falling even further below this curve. Because their study did not include all predictors we used in our work, and therefore we could not calibrate all predictors separately, we applied the same calibration to our scores *p*_del_ = 0.9 · (median score)^6^.

Raw CADD scores were transformed using a sigmoid model *y* = 1*/*(1 + exp(− *A* − *x* + *B*)), with *A* and *B* parameters estimated by Benevenuta et al.^140^. PHRED-scaled scores from FAVOR (protein function annotation PC and conservation annotation PC) were rescaled to the [0, 1] range via 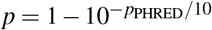 Haploinsufficiency probabilities (*p*_haplo_) were taken directly from the Ensembl VEP annotation.

Because there are no data on indel burden in cardiomyocytes, we estimated *p*_lethal_ for cardiomyocyte indels using the same *p*_lethal | SNVs_*/p*_lethal | indels_ ratio as in brain neurons (similarly to how we estimated indel accumulation rate for cardiomyocytes from the brain data in the Estimating accumulation rates *µ*_0_ of somatic mutation burden section).

After obtaining *p*_lethal SNVs_ and *p*_lethal indels_ for every tissue, we calculated *µ* as:

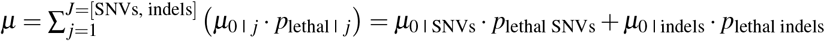

The resulting *p*_lethal_, *µ*_0_, and *µ* for every cell type are summarized in Table 3.

**Table 3.** Mutation accumulation rates and lethality probabilities across cell types used in our modeling.

| Tissue | Cell type | $\mu_0$ for SNVs | $p_{\text{lethal}}$ for SNVs | $\mu_0$ for indels | $p_{\text{lethal}}$ for indels | $\mu$ | $\mu$ (CI low) | $\mu$ (CI high) |
| --- | --- | --- | --- | --- | --- | --- | --- | --- |
| Brain | Neurons | 17.49 | $12.34 \cdot 10^{-5}$ | 6.93 | $3.87 \cdot 10^{-5}$ | $2.44 \cdot 10^{-3}$ | $1.53 \cdot 10^{-3}$ | $3.95 \cdot 10^{-3}$ |
| Heart | Cardiomyocytes | 36.37 | $5.67 \cdot 10^{-5}$ | 14.40 | $3.87 \cdot 10^{-5}$ | $2.61 \cdot 10^{-3}$ | $1.41 \cdot 10^{-3}$ | $4.62 \cdot 10^{-3}$ |
| Liver | Hepatocytes | 52.78 | $2.37 \cdot 10^{-5}$ | 1.16 | $8.74 \cdot 10^{-7}$ | $1.24 \cdot 10^{-3}$ | $0.07 \cdot 10^{-3}$ | $2.25 \cdot 10^{-3}$ |
| Liver | LPCs | 33.73 | $2.37 \cdot 10^{-5}$ | 0.69 | $3.17 \cdot 10^{-8}$ | $0.79 \cdot 10^{-3}$ | $0.43 \cdot 10^{-3}$ | $1.44 \cdot 10^{-3}$ |
| Lung | BBCs | 28.52 | $7.34 \cdot 10^{-5}$ | 2.57 | $5.94 \cdot 10^{-9}$ | $2.08 \cdot 10^{-3}$ | $1.14 \cdot 10^{-3}$ | $3.72 \cdot 10^{-3}$ |

### Estimating other model parameters for each organ

Aside from the mutation accumulation rate *µ*_0_ and the probability of a mutation to be lethal *p*_lethal_, we needed to estimate other parameters to be able to solve our ODEs: organ capacity *K*, critical thresholds of cell population necessary for organ survival *X*_crit_, maximum replication rates *r, r*_*y*_, and *r*_*b*_ (for hepatocytes, LPCs, and BBCs, respectively), Hayflick limit *H*, liver stem capacity *Q*, Hill-type sigmoid parameters *θ* and *n* (for the liver model), and all fractions of different division types that the LPCs and BBCs can undergo.

### Brain modeling parameters

Modeling brain as a whole single organ is difficult, because its regions are rather heterogeneous, with varying energy demands and neuron/non-neuron ratios. As most neurons in the analyzed datasets came from the frontal cortex, we focused on this region to provide the parameters for our model. While there have lots of different estimates of neuron counts in the human brain throughout history, the latest research reports that frontal cortex contains (3.5 ± 0.7) · 10^9^ neurons (40% of brain mass, 34% of brain neurons), with no significant difference between males and females and with no significant decline across the human lifespan^119^.

Defining the critical threshold of cortex neurons necessary for human survival is rather vague: data from functional hemispherectomy (removal of one cerebral hemisphere) in children and adults^141,142^ and frontal lobotomy history^143^ can be used to argue that the frontal cortex is certainly important^144^, but not required for vital autonomic control, suggesting a relatively low survival floor for the *X/K* ratio. However, the same works show that people can survive biologically with far less cortex, but cognition (in the everyday, independent-function sense) fails much earlier — therefore, we switch from “vital for survival” to “vital for cognition” (that is, not demented; capable of independent living). Still, no study gives a neuron-count “cut-point” for being cognitively normal. We inferred the threshold from human data on how much neuronal loss (or closely tracked proxies like cortical thickness/atrophy) accompanies clinically manifest dementia in Alzheimer’s disease (AD) and frontotemporal dementia (FTD). According to those, once ~ 30-50% neuronal loss /marked cortical thinning accrues in the association cortex (including frontal cortex), cognitive deficits are the rule^145–147^; therefore, we place the cognitive “floor” in the middle of this range of neuronal loss, at *X*_crit | brain_ = 0.6*K*.

### Heart modeling parameters

According to the stereological and ^14^C birth-dating studies, the most cited cardiomyocyte (CM) numbers are as follows. The whole heart CM estimates vary widely^148^: some researchers find increase from ~ 10^9^ at birth to ~ 4 · 10^9^ in adults (≤ 20 years)^149^, while others report CM numbers already high at 1 month of age remaining constant through life at (3.2 ± 0.75) · 10^9150^. We treat (3.2 ± 0.75) · 10^9^ as a whole-heart estimate with reasonable uncertainty for adults.

The number of cardiomyocytes is reported to remain constant during the human lifespan^150^, and CM turnover occurs mostly during the first two decades of life; after that, turnover rate declines exponentially in adults from ~ 1%*/*year at the age of 20 years to ~ 0.5%*/*year in the elderly^150,151^. Even though some heart regeneration appears possible^151^, stem cell activity is negligibly minimal, because it is evidently not sufficient to restore heart function after severe injury due to myocardial infarction. Loss of ≥ 40% left ventricle mass (that is, when ~ 0.6*K* remains) was reported to necessitate cardiogenic shock and very high mortality^152^. Even with recent advances in medicine, long-term patient survival is strongly compromised in such severe conditions^153^. Admittedly our model of gradual CM loss should not be matched directly to the cases of acute myocardial infarction, but in case of radical life extension, on the other hand, the remaining cardiac output must be able to sustain human body for a lot longer than the current human lifespan. To account for a possibility of long-term adaptation to the gradually lowering cardiac output, we set critical CM threshold slightly lower than 0.6*K*, at *X*_crit | heart_ = 0.55*K*.

### Liver modeling parameters

#### Liver somatic and stem capacities

We calculated liver hepatocyte capacity via multiplying total liver mass by hepatocellularity (hepatocyte density, in cells per gram). For liver mass, we took estimates from a large autopsy-based study performed by Bell et al.^154^ Regarding hepatocellularity, we used the protein ratio-based measurements by Sohlenius–Sternbeck^155^ estimating (139 ± 25) · 10^6^ hepatocytes/g. LPC definitions vary in terms of namings, morphology, and cellular markers (liver progenitor cells, hepatic stem cells, EpCAM^+^ adult hepatic progenitors, “oval cells”, biliary epithelial cell–derived progenitors, mesenchymal liver stem cells, etc.), introducing notable uncertainty to their population counts. For a mean estimate of *Q* (LPC capacity), we used the most recent measurements from spatial and single-cell transcriptomics data^156^ reporting LPCs as 2.95 ± 1.91% of epithelial liver population. For the lower bound, murine marker-based measurements^157^ reporting LPCs as 0.5% of epithelial liver population can be used. Epithelial liver population comprises LPCs, hepatocytes, and cholangiocytes, therefore we also retrieved hepatocyte (60 ± 10%)^158,159^ and cholangiocyte (4 ± 1%)^160^ fractions of total liver cells to compute *Q*.

Monte-Carlo sampling (500,000 samples) was performed to propagate uncertainty from the respective studies. All parameters were modeled as normal distributions with truncation applied to maintain biological plausibility. The analysis computed total epithelial cells as hepatocytes multiplied by an epithelial multiplier derived from cholangiocyte and hepatocyte fractions, then calculated LPCs as a fraction of this epithelial population. As a result, we obtained the following liver cell capacities:

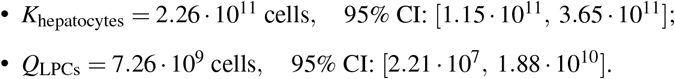

#### Liver critical threshold

Before liver surgery, the future liver remnant (FLR) measure (as determined by computer tomography volumetry) is used as a clinical proxy for how much liver remains after planned resection (partial hepatectomy, also called PHx) — and hence, how many functioning hepatocytes remain. Multiple reviews and clinical practice sources converge on FLR of at least 30% for healthy livers^161,162^. Some meta-reviews even report successful, survivable resections of *>*70% liver volume^163^. Therefore, we estimate the critical threshold for hepatocyte population required for liver survival as *X*_crit | liver_ = 20 *±* 5% of hepatocyte capacity = (0.20 *±* 0.05) · *K*.

#### Hepatocytes proliferation limit

Albeit capable of sustained proliferation *in vivo*, human hepatocytes do not divide as readily *in vitro*, rapidly losing their proliferative ability and function in standard culture and typically surviving only a few divisions before senescence or loss of phenotype^164^. The approaches to culturing murine hepatocytes are also difficult to translate to the human ones^164^, suggesting that the *TERT*^*high*^ hepatocyte subspecies featured in murine livers^97^ might not be as prominent in humans. Excluding the studies of hepatocyte de-differentiation, we can rely on a study which reached at least 40 hepatocyte population doublings^164^ for the lower bound of our proliferation limit: *H*_lower_ = 40. From the upper side, we set the bound at *H*_lower_ = 200 based on the adult human hepatocytes immortalized via telomerase overexpression and inhibition of tumor suppressors^165^. Due to the lack of a better measure, we estimated *H* for hepatocytes via geometric mean: 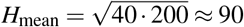 divisions.

#### Hepatocytes replication rate

In healthy human liver (i.e., steady state), diploid hepatocytes replicate on average at *r*_lower_ = 0.71 year^−1166^. To estimate *r*, we leveraged datasets of human liver regeneration after partial hepatectomy with varying resection percentages (that is, varying FLR). During such a short timeframe, we assume that cell mortality is negligible (*µ* → 0) and proliferative potential loss is also negligible (*P/H* → 1). Because we also know from literature that LPCs engage in liver regeneration meaningfully only at 2/3 liver loss and more, the *dX/dt* equation from our Model III becomes very simple:

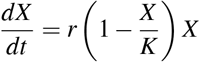

We grouped 9 datasets from 6 different PHx studies based on their *X*_0_*/K* ratio (*X/K* immediately after resection) as follows. Lower bound (LB) group: *X*_0_*/K* = [0.70, 1.00] (less than 1/3 resection, regeneration is not at its fullest)^167–169^; upper bound (UB) group: *X*_0_*/K* = [0.00, 0.35] (large resection encompassing 2/3 PHx and slightly less than that; LPC division contributes to regeneration, so this group cannot be used to give a mean *r* estimate)^170,171^; and a mean estimate (MEAN) group: *X*_0_*/K* = (0.35, 0.70)^168,169,171,172^. We fit *X* (*t*) curves using the equation using the maximum likelihood approach for each dataset propagating the reported CI intervals, and then inferred a combined curve and *r* for each group (Extended Data Fig. 7), which resulted in *r* values and its CI range displayed in Table 5.

**Table 4.** Variables included in the proposed Model II for human brain neurons and heart cardiomyocytes.

| Variable | Description | Brain value and units | Brain source | Heart value and units | Heart source |
| --- | --- | --- | --- | --- | --- |
| $K$ | Somatic cell population capacity | $(3.5 \pm 0.7) \cdot 10^9$ cells<br>(frontal cortex only) | Ref.<sup>119</sup> | $(3.2 \pm 0.75) \cdot 10^9$ cells | Ref.<sup>150</sup> |
| $X_{\text{crit}}$ | Critical population threshold | 0.60K | Refs.<sup>145–147</sup> | 0.55K | Assumed from<br>Refs.<sup>152,153</sup> |

**Table 5.** Hepatocyte replication rates inferred from datasets of liver regeneration after hepatectomy.

| Group | $r \text{ (day}^{-1}\text{)}$ | $r_{\text{CI low}} \text{ (day}^{-1}\text{)}$ | $r_{\text{CI high}} \text{ (day}^{-1}\text{)}$ | $r \text{ (year}^{-1}\text{)}$ | $r_{\text{CI low}} \text{ (year}^{-1}\text{)}$ | $r_{\text{CI high}} \text{ (year}^{-1}\text{)}$ |
| --- | --- | --- | --- | --- | --- | --- |
| LB | $7.81 \cdot 10^{-3}$ | $5.00 \cdot 10^{-3}$ | $12.71 \cdot 10^{-3}$ | 2.85 | 1.83 | 4.64 |
| MEAN | $1.83 \cdot 10^{-2}$ | $1.28 \cdot 10^{-2}$ | $2.74 \cdot 10^{-2}$ | 6.69 | 4.64 | 10.02 |
| UB | 0.158 | 0.114 | 0.205 | 57.87 | 41.75 | 74.76 |

#### Liver progenitor cell replication rate

As mentioned above, the nature of liver stem cells is relatively elusive, and different approaches to defining and quantifying them exist, which makes it challenging to estimate their parameters in our model: maximum stem proliferation rate *r*_*y*_, sigmoid gate parameters *θ*_*X*_, *θ*_*Y*_, *n*_*x*_, and *n*_*y*_, and the three division probabilities *f*_*yy*_, *f*_*xx*_, and *f*_*xy*_. Based on the numerous evidence of LPCs being activated only upon *>* 70% of liver loss^92,95,96^, we assumed both *θ*_*X*_ = 1*/*3 and *θ*_*Y*_ = 1*/*3 (which reflect *X/K* and *Y/Q* ratios, respectively, at which half of LPCs are activated and proliferate).

For *r*_*y*_ estimation, we relied on the upper/lower bounds approach: as upper bound, we inferred *r*_*y* | upper_ from primary-culture population doubling times using:

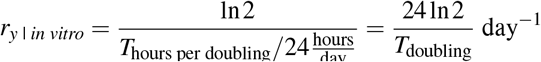

Calculating *r*_*y* | *in vitro*_ from two studies of adult (*T*_*d*_ = 62 h)^173^ and fetal (*T*_*d*_ = 46 h)^174^ human LPCs yields *r*_*y*_ ≈ 0.268 day^−1^ and *r*_*y*_ ≈ 0.362 day^−1^, respectively. Another, later study cultured human hepatocytes-derived LPCs (HepLPCs), biliary epithelium cells-derived LPCs (BecLPCs) and the actual, resident LPCs-derived LPCs (reLPCs) and demonstrated notably shorter doubling times (~ 25.8, ~ 28.5, and ~ 21.6 h, respectively)^175^, which converts to much higher proliferation rates of 0.645, 0.584, and 0.770 day^−1^, respectively. Evidently, even though LPCs do not appear to proliferate in homeostasis (*r*_*y* | lower_ → 0), they are still capable of a much higher activity in a wide range of conditions. We provide an estimate for *r*_*y*_ via a geometric mean of adult LPC-derived upper bounds and an approximation for *r*_*s* | lower_ ≈ 10^−4^ day^−1^ ≈ 0.037 year^−1^:

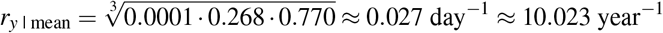

This value is still lower than the lowest *in vitro*-inferred upper bound (0.268 day^−1^), and is not pegged down to zero, as it would be if we considered *r*_*y*_ only at homeostasis.

#### Liver progenitor cell division probabilities

The proportions between division types change relatively to the current deficits of both stem (*Y/Q*) and somatic (*X/K*) cell populations. In homeostasis, asymmetric divisions (*f*_*xs*_) predominate, and the symmetric self-renewal (*f*_*yy*_) and differentiation (*f*_*xx*_) cancel each other out, according to the neutral-drift models^176^. After a severe injury causing widespread cell loss, stem cells self-renew rapidly in order to later switch to symmetric differentiation and quickly replenish the organ’s functional capacity^177,178^. Therefore, all division type probabilities should be modeled as functions of both *y* = *Y/Q* and *x* = *X/K*. At any *y* or *x*, the division fractions must be non-negative and sum to one: *f*_*xx*_ + *f*_*yy*_ + *f*_*xy*_ = 1.

A convenient choice to model such relationships is to form utilities for each division outcome that are linear (or affine) functions of *y* and *x*, and then convert these utilities to probabilities by a softmax^179^. The softmax approach enforces positivity and the sum-to-one automatically. Because the softmax depends only on utility differences, one of the utilities can be fixed as a reference. We choose *U*_*xy*_ = 0 as the baseline (equivalently we could subtract the same constant from all utilities). This zero baseline allows us to avoid an extra set of parameters that would otherwise be redundant and ensures identifiability: without fixing one of the utilities, the parameters are under-determined. Thus, the system of utilities can be parametrized as:

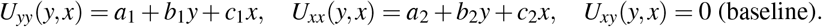

From these utilities, we can derive the probabilities of each division type as:

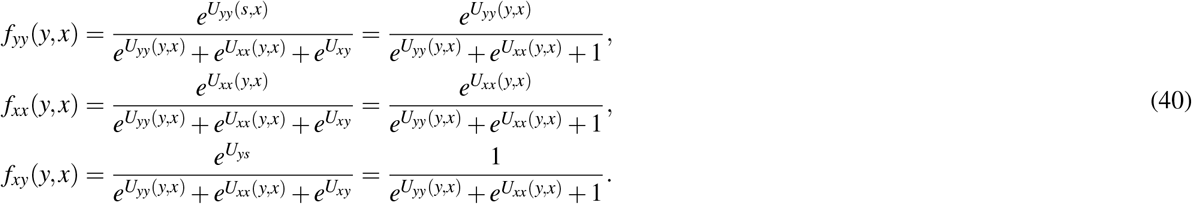

As one can infer 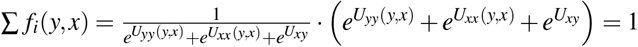 which supports the sum-to-one rule. The softmax lets us transform observed probabilities into log-odds (utilities) very simply. Divide each observed probability by the baseline *f*_*xy*_ and take the natural logarithm:

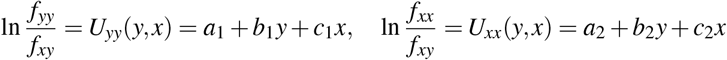

For each state (*y, x*) and triplet (*f*_*yy*_, *f*_*xx*_, *f*_*xy*_) observed at this state, we thus have two linear equations as the ones above. Each parameter has a simple interpretation:

- *a*_1_, *a*_2_ — baseline utilities (intercepts) for *U*_*yy*_, *U*_*xx*_ when *y* = 0, *x* = 0.
- *b*_1_ — how self-renewal utility changes with stem pool *y*. If *b*_1_ *<* 0, higher stem occupancy reduces the drive to self-renew.
- *b*_2_ — how differentiation utility changes with *y*. If *b*_2_ *>* 0, higher stem occupancy increases tendency to differentiate.
- *c*_1_, *c*_2_ — how each utility changes with the somatic occupancy *x*. Typical sign patterns: higher *x* (full somatic pool) should suppress differentiation (negative *c*_2_) and suppress the need for self-renewal (negative *c*_1_), but the exact fitted signs follow the data.

Because utilities enter the probabilities exponentially, a change of 1 unit in a utility multiplies the corresponding odds by *e*^1^ ≈ 2.7 — so magnitudes are interpretable in odds terms.

Given several anchor states (*y, x*) with the respective (*f*_*yy*_, *f*_*xx*_, *f*_*xy*_) triplets, we can solve the softmax system algebraically. The problem with LPCs, however, is that the precise division fractions are hard to find in the literature. One study^177^ quantified symmetric self-renewals and asymmetric divisions of Lgr5^+^ stem cells in murine livers injured via toxin administration (Lgr5^+^ supposedly marks damage-induced LPCs)^92^. The injured mice were treated with either FXR (farnesoid X receptor) agonist to force LPC symmetric self-renewal, or PPAR*α* (peroxisome proliferator-activated receptor *α*) agonist to promote asymmetric divisions. After FXR treatment, the probabilities were demonstrated as *f*_asymmetric_ = *f*_*xy*_ ≈ 15%, *f*_symmetric_ ≈ 85%. After PPAR*α* treatment: *f*_asymmetric_ = *f*_*xy*_ ≈ 80%, *f*_symmetric_ ≈ 20%.

Because PPAR*α* activation drives LPCs to quiescence and maintenance, we assumed that the PPAR*α*-induced fractions can be used to approximate division probabilities at homeostasis. We decomposed the fraction of symmetric divisions into *f*_*yy* | homeo_ = *f*_*xx* | homeo_ = 1*/*2 · *f*_symmetric | PPAR*α*_ ≈ 20*/*2 = 10%, so that self-renewal and differentiation cancel each other at homeostasis, leaving most divisions asymmetric *f*_*xy* | homeo_ = 80%.

On the contrary, the FXR-induced triplet corresponds to the state of hyperactive self-renewal following injury. In physiological injury response without exogenous FXR activation, differentiation would not be impeded, and a fraction of symmetric divisions would continue leading to it: we assume it to be the same as homeostasis, *f*_*xx* | stem wave_ = *f*_*xx* | homeo_ = 10%; then *f*_*yy* | stem wave_ = *f*_symmetric | FXR_ − *f*_*xx* | stem wave_ = 85 − 10 = 75%. As this study did not trace the later differentiation of FXR-induced *Lgr5*^*+*^ daughters, we can only assume that symmetric differentiation rises to the same levels as does symmetric self-renewal in order to supply functional hepatocytes to the injured organ from the newly made LPCs: *f*_symmetric | stem wave_ = *f*_symmetric | differentiation wave_. Hence, swapping the two kinds of symmetric divisions betwen each other, *f*_*xx* | differentiation wave_ = *f*_*yy* | stem wave_ = 75% and *f*_*yy* | differentiation wave_ = *f*_*xx* | stem wave_ = 10%. As the exact (*y, x*) pairs for stem and differentiation waves are not known, we hypothesized these pairs based on our addition-based sigmoid rule for stem cell replication (29) and (30), according to which LPC proliferation rate *ρ*_*y*_ is high at *y* = 1*/*3, at *x* = 1*/*3, and continues to be high at *y* = 2*/*3 if *x* is still 1*/*3.

Thus, the three calibration points become as shown in Table 6.

**Table 6.** LPC division probabilities inferred from Ref.^177^.

| State | y | x | $f_{yy}$ | $f_{xx}$ | $f_{xy}$ |
| --- | --- | --- | --- | --- | --- |
| Homeostasis | 1 | 1 | 0.10 | 0.10 | 0.80 |
| Injury, stem wave | 1/3 | 1/3 | 0.75 | 0.10 | 0.15 |
| Injury, differentiation wave | 2/3 | 1/3 | 0.10 | 0.75 | 0.15 |

Solving the softmax system at these calibration points with the `lstsq` function of *numpy* Python package, we obtained:

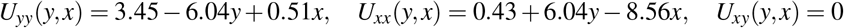

These utilities are then fed in the softmax system (40) to compute LPC division probabilities at each *x* and *y* of our modeling (Extended Data Fig. 9).

#### Liver modeling summary

The resulting values for all liver-related variables included in our modeling are summarized in Table 7.

**Table 7.** Variables included in the proposed Models IIIA and IIIB for human hepatocytes and LPCs.

| Variable | Description | Value and units | Source |
| --- | --- | --- | --- |
| $K$ | Somatic cell population capacity | $2.26 \cdot 10^{11}$ cells | Computed from Refs.<sup>154, 155</sup> |
| $Q$ | Stem cell population capacity | $7.26 \cdot 10^9$ cells | Computed from Refs.<sup>156, 158–160</sup> |
| $X_{\text{crit}}$ | Critical population threshold | $0.20K$ | Assumed from Refs.<sup>161–163</sup> |
| $r$ | Maximum somatic replication rate | $7.45 \text{ year}^{-1}$ (LB: 1.79, UB: 75.32) | Computed from Refs.<sup>167–172</sup> |
| $H$ | Proliferation limit | 90 divisions (LB: 40, UB: 200) | Computed from Refs.<sup>164, 165</sup> |
| $r_y$ | Maximum stem replication rate | $10.0 \text{ year}^{-1}$ (LB: 0.037, UB: 281.3) | Computed from Refs.<sup>173–175</sup> |
| $f_{yy}$ | Symmetric self-renewal probability | Inferred as $f_{yy}(y, x)$ | Computed from Ref.<sup>177</sup> |
| $f_{xx}$ | Symmetric differentiation probability | Inferred as $f_{xx}(y, x)$ | Computed from Ref.<sup>177</sup> |
| $f_{xy}$ | Asymmetric division probability | Inferred as $f_{xy}(y, x)$ | Computed from Ref.<sup>177</sup> |

### Lung modeling parameters

#### Basal cell capacity

The data on somatic mutagenesis in lungs came from the proximal bronchial basal cells; however, it is unreasonable to estimate basal cell capacity based only on the proximal bronchi: the airway tree is continuous, and epithelial cells can proliferate along its whole length. Therefore, to estimate basal cell capacity *K* in human airway epithelium, we employed the measurements of tracheal geometry and histology data of the whole human tracheobronchial tree. The inputs for this estimation are: CT-derived tracheal lumen volume *V* = 32.0 *±* 8.3cm^3^ and tracheal length *L* = 102.8 *±* 9.9 mm^180^, diameter reduction factors *k* = 0.79 and 0.94 (depending on generation number) to generate bronchial generation-wise diameters via a symmetric Weibel model^181,182^, a length-to-diameter ratio LtD = 1.46 ± 0.15 to convert diameters into bronchial generation lengths^181^, total conducting airway surface area *A*_measured_ = 2, 471 ± 320 cm^2183^, total epithelial cell count *N*_epithelial_ = 10.5 · 10^9183^, number of conducting airway generations *G* = 25^182^, basal cell fractions at large and terminal airways^184^, and sex-specific total lung volumes (TLVs)^185^.

Each bronchial generation has *n*_*g*_ = 2^*g*^ airways. This is an idealization: asymmetric branching will change counts and therefore area allocation, but 2^*g*^ is the standard Weibel-type starting point. Trachea is normally oval-like, but we approximate it by computing the equivalent circular tracheal diameter *D*_0_ from its volume *V* and length *L* as

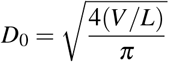

Diameter change by each bronchial generation *g* depends on the per-generation reduction factor *k*, which changes from 0.79 before ~ 15th generation to 0.94 after it^181^. We used the following piecewise model, which ensures *D*_*g*_ is continuous at the transition (no artificial jump):

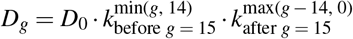

Given a generation-specific length-to-diameter ratio LtD = *L*_*g*_*/D*_*g*_ (dimensionless), the lateral surface area of one bronchial branch in generation *g* would be

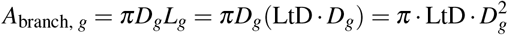

Each generation’s area is 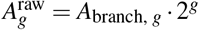. These areas are relative predictions based on idealized assumptions (perfect symmetry, cylinders, no irregularities, etc.). Morphometry^183^ reports a measured total conducting airway surface area *A*_measured_ and a total epithelial cell count *N*_epithelial_, so we enforce the empiric total area via the scaling factor

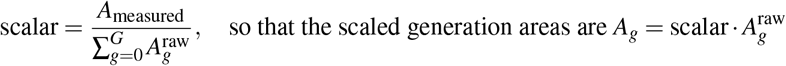

which preserves the model’s relative distribution of areas while ensuring the sum of modeled areas equals the empiric total. We treat *N*_epithelial_ as a reference at TLV_combined sexes_. For each sex group with TLV equal to TLV_sex group_, we scale the epithelial count linearly:

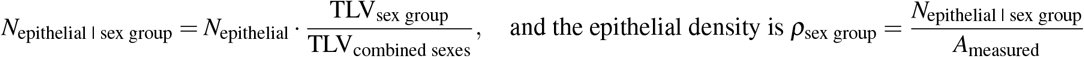

Basal cells are primarily located in the proximal conducting airways (trachea, bronchi, and bronchioles)^186^. In trachea, basal cells make up roughly 40 ± 5% of the epithelium (at least in mice)^106^. Normal human bronchial epithelium contains 6–31% basal cells, with proportion varying along the proximal-distal axis^184^. The basal cell-containing conducting epithelium extends distally to terminal bronchioles of about 0.5 mm diameter and only the respiratory bronchioles are lined by a simple cuboidal epithelium lacking basal cells^187^. We used linear interpolation to calculate the basal fractions for *G* = 25 bronchial generations (between the 31 and 6% basal fraction endpoints): let *p*_*g*_ be the fraction of epithelial cells at generation *g* that are basal. For each *g*-th generation,

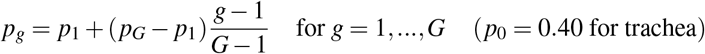

Each generation contains *K*_*g*_ = *A*_*g*_ · *ρ*_sex group_ · *p*_*g*_ basal cells. Finally, the total basal cell capacity *K* is

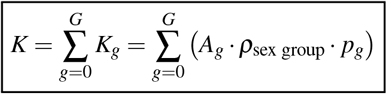

Uncertainty is propagated by Monte-Carlo sampling (500,000 draws): all uncertain inputs (tracheal *V, L*, LtD, *N*_epithelial_, TLVs, and class basal fractions *p*_*g*_) are drawn from truncated normal distributions to avoid nonphysical negative draws from the normal distributions (implemented with `truncnorm` from the *scipy* Python package).

As a result, we obtained the following airway basal cell capacities:

- *K*_hepatocytes_ = 1.03 · 10^9^ cells, 95% CI: [5.73 · 10^8^, 1.59 · 10^9^]

#### Lung critical threshold

Estimating the critical threshold for the *B/K* ratio is difficult, because there is no direct human experiment that reports a single numeric threshold. From the transplantation studies, we can assume that at least 60 million basal cells are required to restore lung function^188^, placing the lower bound at 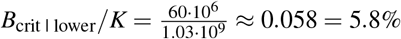. From lung resection studies showing that the net loss of at least 10% lung volume is perfectly tolerable^189,190^, we can assume that the same proportion of basal cell loss is also tolerable, therefore *B*_crit | upper_*/K* = 0.90 = 90%. We estimated *B*_crit_ as a geometric mean of these bounds, making 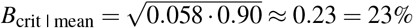.

#### Basal cell replication rate

To estimate maximum replication rate for BBCs, we again resorted to the upper/lower bound logic. A steady-state study of murine tracheal basal cells by Watson et al.^106^ found that these cells divide, on average, once per 11 ± 4.4 days. From it, we can infer the lower bound for BBC replication rate (assuming tracheal and bronchial basal cells have similar cycling dynamics) of:

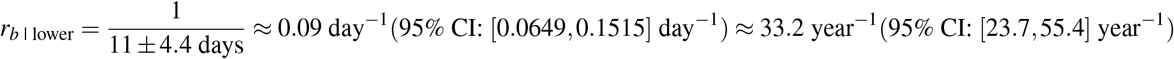

In two studies by Peters-Hall et al.^104,105^, BBCs were cultured in different conditions, exploring their maximum number of population doublings (PD): at 21% or 2% O_2_, with or without Rho-associated protein kinase inhibition (ROCKi), in normal and cystic fibrosis (CF)-derived BBCs. We assumed that under extremely mild, stress-free conditions (2% O_2_, ROCKi, healthy donor), BBCs show virtually unrestricted proliferative potential (with no differentiation bias). Hence, the settings explored by Peters-Hall et al. can approximate the maximum replication rate.

Both studies feature population doubling curves *PD*(*t*). To estimate *r*_*b* | upper_, recall that, under unrestricted growth:

*PD*(*t*) = number of doublings needed to get from *B*_0_ to *B*(*t*)

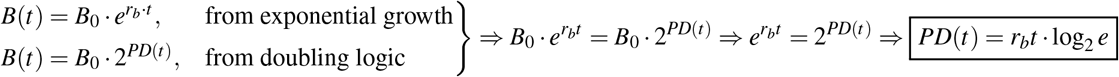

Thus, solving for *r*_*b*_:

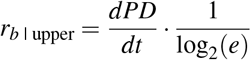

After fitting a linear regression on data points (corresponding to 2% O_2_, ROCKi, healthy donors) provided in Figure 2A of Ref.^104^ and Figure 1A of Ref.^105^, we obtained slopes of ~0.83 and ~0.41 BBC population doublings (PD) per day (*dPD/dt*), respectively (Extended Data Fig. 11). Estimating *r*_*b* | upper_ from each of those yields 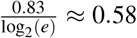 day^−1^ and 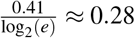 day^−1^. To find *r*_*b* | mean_ and its 95% CI, we calculated geometric mean between the one lower and two upper bounds and propagated the uncertainty from *r*_*b* | lower_ measurement error and *r*_*b* | upper_ fitting errors (via Monte-Carlo sampling with *N* = 10, 000 samples), yielding:

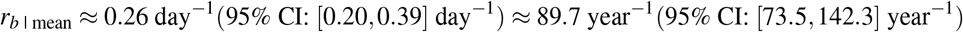

#### Basal cell proliferation limit

In standard culturing conditions, airway basal cells proliferate for 30-40 PDs^104,191^, but the 21% oxygen concentration in these conditions is not physiological: basal cells do not come in direct contact with ambient atmospheric oxygen in the living airways. To mitigate the effect of oxidative damage on BBC replication limits, the ROCKi studies used 2% O_2_ to reach *>*220 PDs^104,105^, but they did so without reporting the exact O_2_ concentration of the *in vivo* BBC niche. Other studies incured hypoxia response in basal cells at *<*1%^192^ and, to some extent, at 5% O_2_ (considered normoxic for many other tissues)^193^, suggesting that the 2% conditions are sub-physiological. To estimate BBC replication limit, we set *H*_lower_ = 40 from the 21% O_2_ conditions, *H*_upper_ = 220 from the 2% O_2_ conditions, and perform linear interpolation between these two endpoints, yielding *H*_mean_ ≈ 172.6 ≈ 170 divisions at 7% O_2_ (putative oxygen concentration at the BBC level; slightly above the still hypoxic 5% O_2_).

#### Basal cell division probabilities

In contrast to the liver progenitor division probabilities, we cannot derive BBC division probabilities as functions of both *b* = *B/K* and some *x* = *X/*(differentiated cell capacity), because there are no data on somatic mutagenesis in the differentiated daughters of basal cells yet, and, therefore, we cannot consider them in our model. Hence, we derive the division fractions only as functions of *b*. We also have to assume that *f*_*bb*_ = *f*_*xx*_ at any *b* (assuming differentiation and self-renewal cancel out in the long run), because there is no *x* dimension which we relied on to model the *f*_*bb*_ and *f*_*xx*_ dynamics in case of liver progenitors.

The aforementioned study of murine tracheal basal cells at homeostasis^106^ reports the following division fractions: *f*_*xb* | homeo_ = 0.94 ± 0.03 = 94 ± 3%, *f*_*bb* | homeo_ = *f*_*xx* | homeo_ = 0.3 = 3%. In another study, influenza infection was shown to shift basal cell division to *f*_symmetric | injury_ = 0.48 = 48%, thus *f*_asymmetric | injury_ = *f*_*xb*_ = 1 − 0.48 = 0.52 = 52%^194^. From our *f*_*bb*_ = *f*_*xx*_ assumption, *f*_*bb* | injury_ = *f*_*xx* | injury_ = 0.48*/*2 = 0.24 = 24%. This 8-days long influenza infection resulted in the decrease of ciliated area per micron from ~ 4.2 to ~ 0.5 and in the decrease of Foxj^+^ (ciliated airway cell marker) cells per micron from ~ 0.024 to ~ 0.005 in infected murine airways^194^, suggesting that the population of differentiated ciliated cells dropped to levels somewhere between 0.5*/*4.2 ≈ 0.12 = 12% and 0.005*/*0.024 ≈ 0.21 = 21%. Assuming that the population of basal cells experienced a similar drop, we set the *B/K* anchor at injury to 25% (or 1/4).

We leveraged these mouse-based values as the best existing estimate of airway basal cell division fractions at homeostasis and during regeneration and constructed the softmax system, resembling our approach to deriving the LPC division probabilities. Let *b* = *B/K*. Similarly to the LPCs, the system of utilities can be parametrized as:

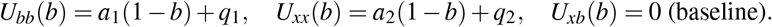

Because of our *f*_*bb*_ = *f*_*xx*_ assumption, *a*_1_ = *a*_2_ = *a* and *q*_1_ = *q*_2_ = *q*. With *a >* 0, decreasing *b* increases (1 − *b*) and thus increases *U*_*bb*_(*b*) = *U*_*xx*_(*b*).

Solving this softmax system for the anchors at injury and homeostasis as before, we obtain *a* ≈ 3.56, *q* ≈ − 3.45, and therefore:

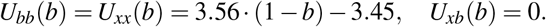

Thus, we compute BBC division probabilities at each *b* of our modeling as (Extended Data Fig. 12):

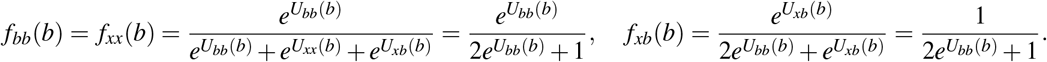

#### Lung modeling summary

The resulting values for all lung-related variables included in our modeling are summarized in Table 8.

**Table 8.** Variables included in the proposed Model IV for human airway basal cells.

| Variable | Description | Value and units | Source |
| --- | --- | --- | --- |
| $K$ | Basal cell population capacity | $1.03 \cdot 10^9$ cells | Computed from Refs.<sup>180–185</sup> |
| $B_{\text{crit}}$ | Critical population threshold | $0.23K$ (LB: 0.058, UB: 0.90) | Computed from Refs.<sup>188–190</sup> |
| $r_b$ | Maximum basal cell replication rate | $0.26 \text{ day}^{-1}$ | Computed from Refs.<sup>104–106</sup> |
| $H$ | Proliferation limit | 170 divisions (LB: 40, UB: 220) | Computed from Refs.<sup>104, 105, 191–193</sup> |
| $f_{bb}$ | Symmetric self-renewal probability | Inferred as $f_{bb}(b)$ | Computed from Refs.<sup>106, 194</sup> |
| $f_{xx}$ | Symmetric differentiation probability | Inferred as $f_{xx}(b)$ | Computed from Refs.<sup>106, 194</sup> |
| $f_{xb}$ | Asymmetric division probability | Inferred as $f_{xb}(b)$ | Computed from Refs.<sup>106, 194</sup> |

### Fréchet-Hoeffding bounds

To quantify the uncertainty in the joint survival function of multiple ograns due to unknown dependence among organ failure times, we employed the Fréchet–Hoeffding bounds — sharp extremal limits on any joint distribution with given marginals^74^.

For a joint survival function *S* (*t*, …, *t*) with marginal survival functions *S*_*i*_(*t*) = ℙ(*T*_*i*_ *> t*), the Fréchet–Hoeffding bounds are:

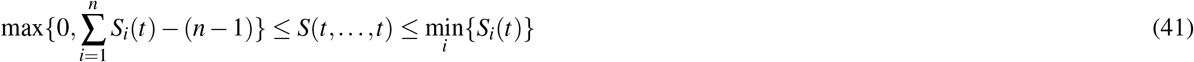

Note that these bounds apply to the joint survival at a common time point *t* (i.e., *t*_1_ = *t*_2_ = … = *t*_*n*_ = *t*), which is relevant when estimating organismal survival as a time until any organ fails. The upper bound corresponds to comonotonicity (perfect positive dependence), where all organs fail simultaneously; the lower bound corresponds to the most antagonistic dependence compatible with the given marginals. Notably, while the upper Fréchet–Hoeffding bound is always attainable (via comonotonicity), the lower bound is sharp only for *n* = 2. For *n* ≥ 3 this bound remains valid but may not be achievable by any joint survival distribution with the given marginals — i.e., it is not necessarily tight. Nevertheless, it is the best possible universal lower bound that depends solely on the marginal survival functions^195^.

In our case, the number of marginals is 4, which means that the inequalities change to:

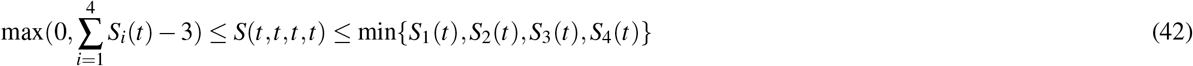

As a reference model under the assumption of mutual independence, we use:

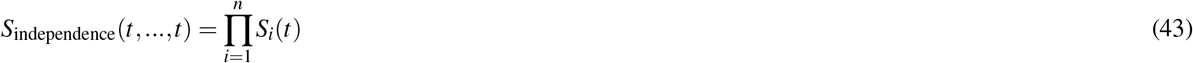

Finally, to account for the background (non-organ-specific) mortality, we multiply the resulting joint survival function and its bounds by an exponential baseline survival term 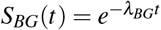 yielding the adjusted survival estimates used in lifespan prediction.

### Monte-Carlo simulation for empirical survival probability attribution

Because Model III variants provide intractable survival probability, we estimate it using Kaplan-Meier. Failure dynamics were governed by a mechanistic ordinary differential equation (ODE) system describing the interactions between state variables.

To account for uncertainty in biological parameters (e.g., mutation rates, carrying capacities, regenerative rates), we performed Monte Carlo simulations under a log-normal parameterization (justified by multiplicative biological noise). For each simulation:

1. Parameters were sampled from a log-normal distribution fitted to empirical means and standard error derived from the 95% confidence intervals.
2. The ODE was integrated forward in time until *X* (*t*) crossed *X*_crit_ or reached maximum time of a simulation *t*_max_ = 10^5^ years, thus incorporating right censoring.
3. Death times were estimated from 10^6^ independent simulations forming a synthetic cohort.

The survival function *S*(*t*) = ℙ(*T > t*) was estimated non-parametrically using the Kaplan–Meier (KM) estimator, with right-censored observations at *t*_max_. Confidence intervals for the KM curve were computed using the Greenwood’s formula^196^. Additionally, to model background (non-organ-specific) mortality, the mechanistic survival estimate was multiplicatively combined with an exponential random mortality term 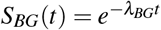, where *λ*_*BG*_ was estimated from human demographic data.

Note that our simulation of the synthetic population uses samples from the parameters that are estimated for the mean distribution.

### Hazard rate estimation

#### Model II

For the brain and heart models, we derived a semi-analytical expression for the organ survival probability:

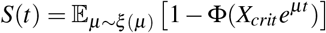

where Φ(…) is the cumulative distribution function (CDF) of the organ capacity *K*.

Using the identity 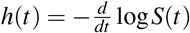, we obtain the corresponding hazard rate:

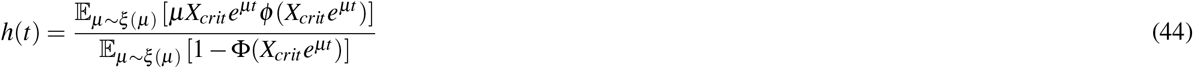

where ϕ (…) is the probability density function (PDF) of *K*.

To estimate the expectation over the distribution *ξ* (*µ*), we used the Monte-Carlo method with 10,000 samples.

#### Model III

For the liver and lungs, where a closed-form solution for the survival probability is not available, we employed the nonparametric Nelson-Aalen estimator for the cumulative hazard function Λ(*t*):

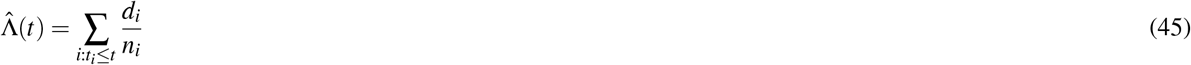

where *t*_*i*_ are the observed event times, *d*_*i*_ is the number of failures at time *t*_*i*_ and *n*_*i*_ is the amount of subjects at risk just prior to *t*_*i*_.

To estimate the hazard rate *h*(*t*) from the cumulative hazard, we differentiated 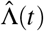 numerically and applied a spline smoothing function from `scipy.interpolate.UnivariateSpline` with the power *k* = 5. This helps to reduce noise and obtain a smooth hazard rate curve.

## Acknowledgements

This study was supported by the Russian Science Foundation [25-71-20017 to E.K.].

## Author contributions statement

E.E.* and V.F.* designed the dynamical system. E.E.* collected data, estimated model parameters, created plots and supplementary figures, and wrote the manuscript. V.F.* designed the distribution-based modeling of population survival, wrote the manuscript, and prepared the project’s GitHub page. L.M.* collected data, wrote the manuscript, participated in the dynamical system design, and created the artwork. E.E.K. supervised the study. D.K. conceived and supervised the study, wrote the manuscript, and collected data. All authors reviewed the manuscript.

* These authors contributed equally to this work.

## Data and code availability

Data and notebooks including the model and parameters calculation, which are necessary to reproduce the results of this paper, are provided in the GitHub repository at https://github.com/ComputationalAgingLab/somatic-mutations-simulation.

## Extended Data Figures

**Extended Data Figure 1.**
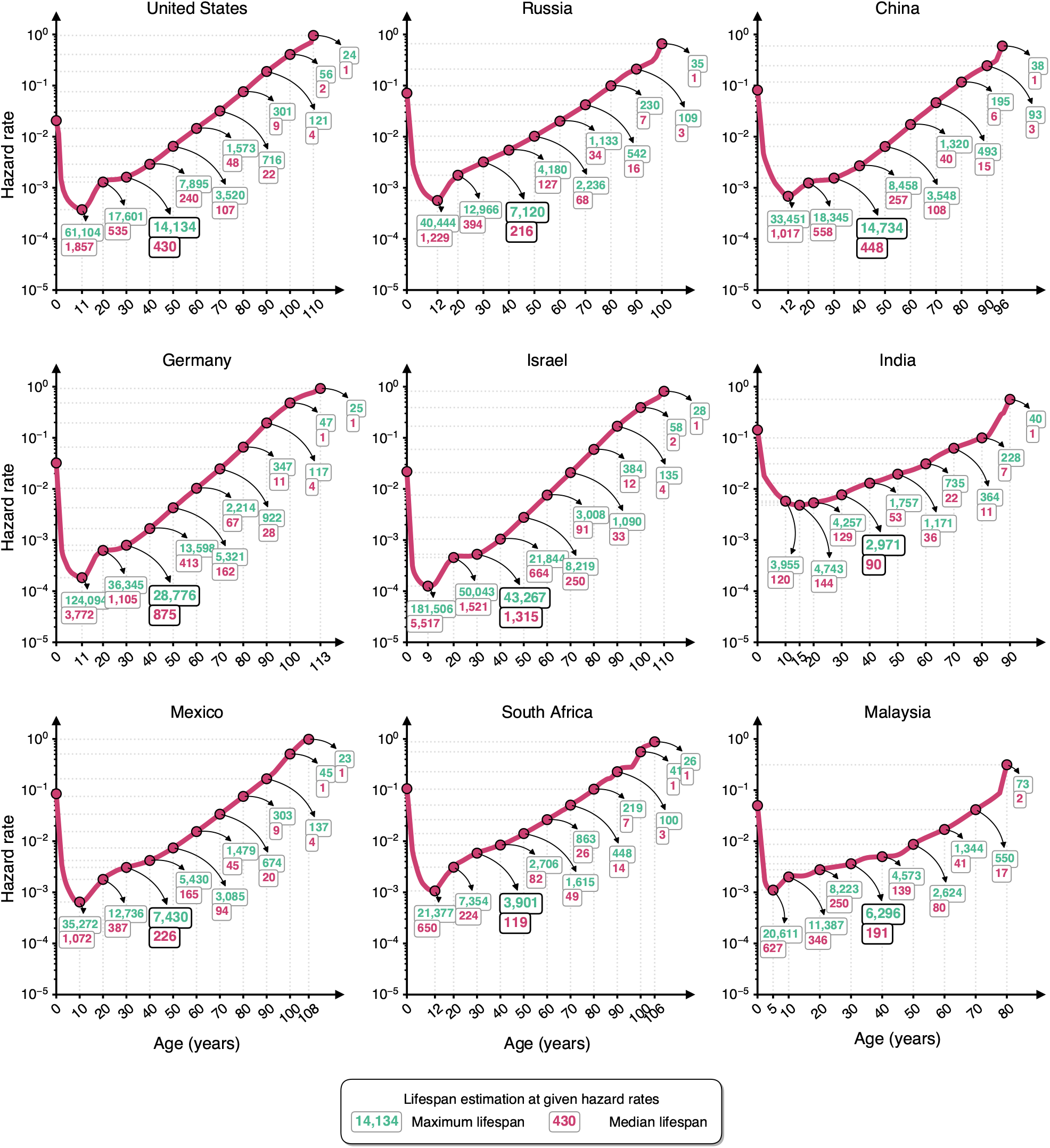
Proposed incremental framework for modeling organismal aging. **a** Among the hallmarks of aging, we focus on genomic instability which is particularly challenging to reverse. **b** Somatic mutations accumulate throughout the human lifespan and correlate strongly with chronological age. **c** Normal survival curve reaches zero at the maximum documented human lifespan of 122 years. Here we ask: *What would the maximum achievable lifespan be if somatic mutations were the sole driver of aging?* It would presumably be more than if all drivers of aging are considered (“Normal aging”) and less than in case of “No aging” (no mortality increase over time). **d** The human body can be modeled as a complex reliability system^56,57^ composed of parallel and series elements; failure of any critical element leads to death. **e** Schematic representation of the hepatocyte population dynamics: starting at full capacity (100%), cell population declines over time, and organ failure occurs upon reaching a certain critical threshold which corresponds to the minimally viable population size. **f** Our incremental modeling framework: three successive stages of model refinement, each introducing an additional layer of biological complexity. Model I considers constant hazard rate (*λ*) due to background mortality fixed at some age. Model II adds mortality due to somatic mutations to overall mortality. Model III accounts for cell replication, which is supposed to compensate partially for cell loss.

**Extended Data Figure 2.**
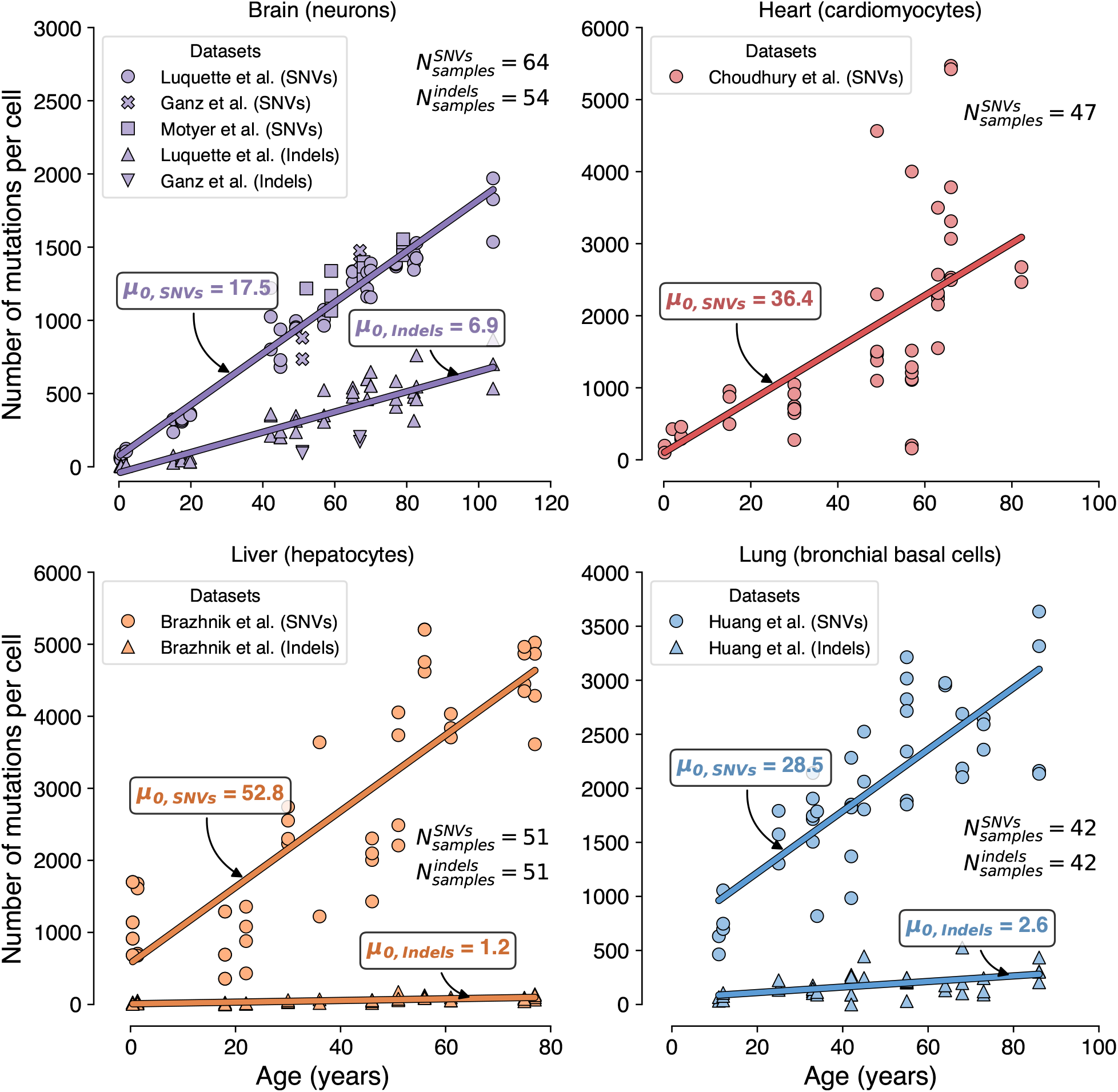
Results of the incremental modeling framework for predicting maximum human lifespan under somatic mutation accumulation. **a** Model I: baseline constant-risk model which accounts only for background (BG) mortality *λ*_BG_. **b** Hazard rate (HR) estimated for different ages in the US population. According to Model I, maximum and median lifespans are defined as ages at which the survival probability drops to 1*/N* (where *N* is the global human population) and to 0.5 (or 50%), respectively. Both lifespans are calculated using HR frozen at every 10 years, except for the point of lowest mortality (HR freeze at 11 years instead of 10). The lifespans calculated for the 30-years mortality are highlighted with black frames and larger font size. **c** Simulated survival curves for Model I across different ages of HR freeze. Age numbers in the legend correspond to the numbers along the curves. The curve calculated for the 30-years mortality is highlighted with a black frame around its age number. **d** Model II: adds organ-specific somatic mutation accumulation which causes cell death at rate *µ* as an additional source of mortality risk *λ*_mut_. **e** Cell population dynamics simulated according to Model II for brain (left) and heart (right). Organ failure occurs when cell population falls below a critical threshold *X*_crit_. **f** Survival curves simulated according to Model II for brain (left) and heart (right) under three scenarios: BG-only, organ-only, and combined (mortality from organ-specific accumulation of somatic mutations and BG mortality). **g** Model IIIA: further includes cell replication via function *ρ*(P), where P denotes the average replication potential of the whole cell population. **h** Somatic cell population dynamics simulated under Model IIIA for hepatocytes. **i** Survival curves simulated for Model IIIA. **j** Model IIIB: extends IIIA by adding adult stem cells (including stem cell proliferation *ρ*_*y*_ with unlimited potential) which replenish the somatic pool at rate *σ* and are also prone to death from somatic mutations at rate *µ*_*y*_. **k** Somatic hepatocyte population dynamics simulated according to Model IIIB in the presence of liver progenitor cells (LPCs); the death threshold is not reached within 100,000 years of simulation. **l** Survival curves simulated for Model IIIB. **m** Model IIIC: stem-like cell type which exhibits mutation-driven mortality (at rate *µ*_*b*_), differentiation, and limited proliferation (at rate *ρ*_*b*_ = *f* (*P*)). **n** Lung cell population dynamics simulated under Model IIIC for bronchial basal cells. **o** Survival curves simulated for Model IIIC. In panels **e, h, k**, and **n**, shaded regions represent percentile bands (from the 2.5th to the 97.5th in 5% increments) for the simulated *X* (*t*) trajectories; bright solid line corresponds to the median trajectory; red dotted line is the critical threshold of cell population required for organ survival (*X*_crit_); the time at which the median cell population trajectory crosses this critical threshold is annotated as *t*_crit_. In panels **f, i, l**, and **o**, horizontal dotted gray lines correspond to survival probabilities of 0.5 and 1*/N*; median and maximum lifespans are annotated for organ-only and combined survival curves (and for the BG curve in **f**) as *t*_50%_ and *t*_1*/N*_, respectively. In panels **i, l**, and **o**, survival curves for models calculated using BG and combined mortalities overlap each other and are represented next to each other for visualization purposes only.

**Extended Data Figure 3.**
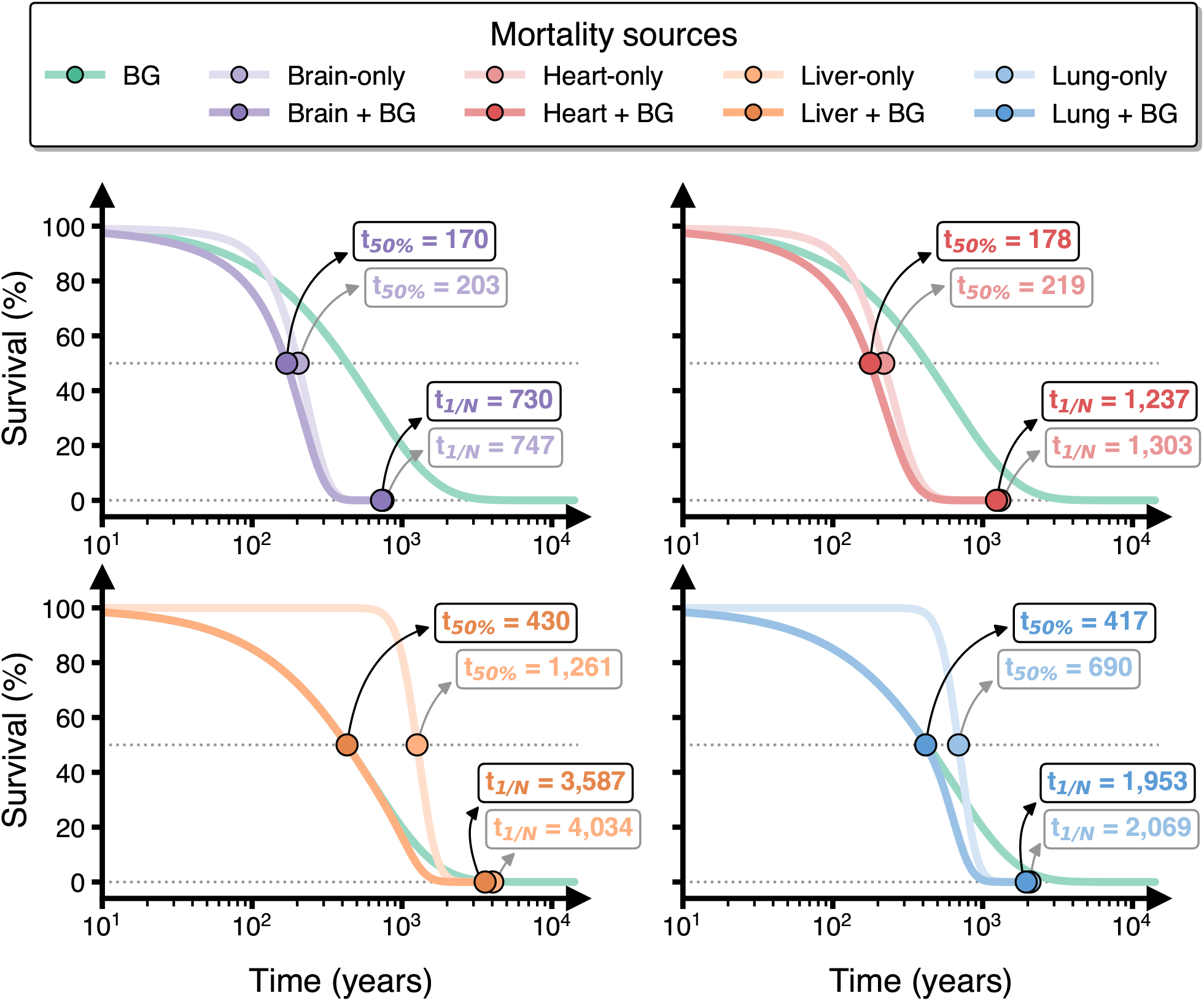
Multi-organ integration reveals theoretical lifespan limits under somatic mutation-driven aging. **a** The organism is modeled as a series reliability system in which four critical organs (brain neurons, heart cardiomyocytes, liver hepatocytes, and respiratory basal epithelium) age independently through somatic mutation-driven cell loss, alongside baseline mortality (*λ*_BG_) fixed at age 30 levels. Failure of any single organ causes organismal death. Under the independence assumption, overall survival is the product of individual organ survival functions. **b** Population survival curves under different inter-organ dependency assumptions. The independence model is represented by solid black line. Fréchet bounds establish theoretical limits shown as dashed gray lines: from perfect positive dependence (upper bound) assuming that all organs age in synchrony, to perfect negative dependence (lower bound) assuming anti-correlated aging. Shaded region with diagonal hatches encompasses all mathematically possible dependency structures. Pink line indicates normal aging (for the US population). Gray horizontal dotted lines correspond to survival probabilities of 0.5 and 1*/N*. Median and maximum lifespans are annotated as *t*_50%_ and *t*_1*/N*_, respectively, for the multi-organ and normal aging models. Survival curves for models IIIB and IIIC overlap each other and are represented next to each other for visualization purposes only.

**Extended Data Figure 4.**
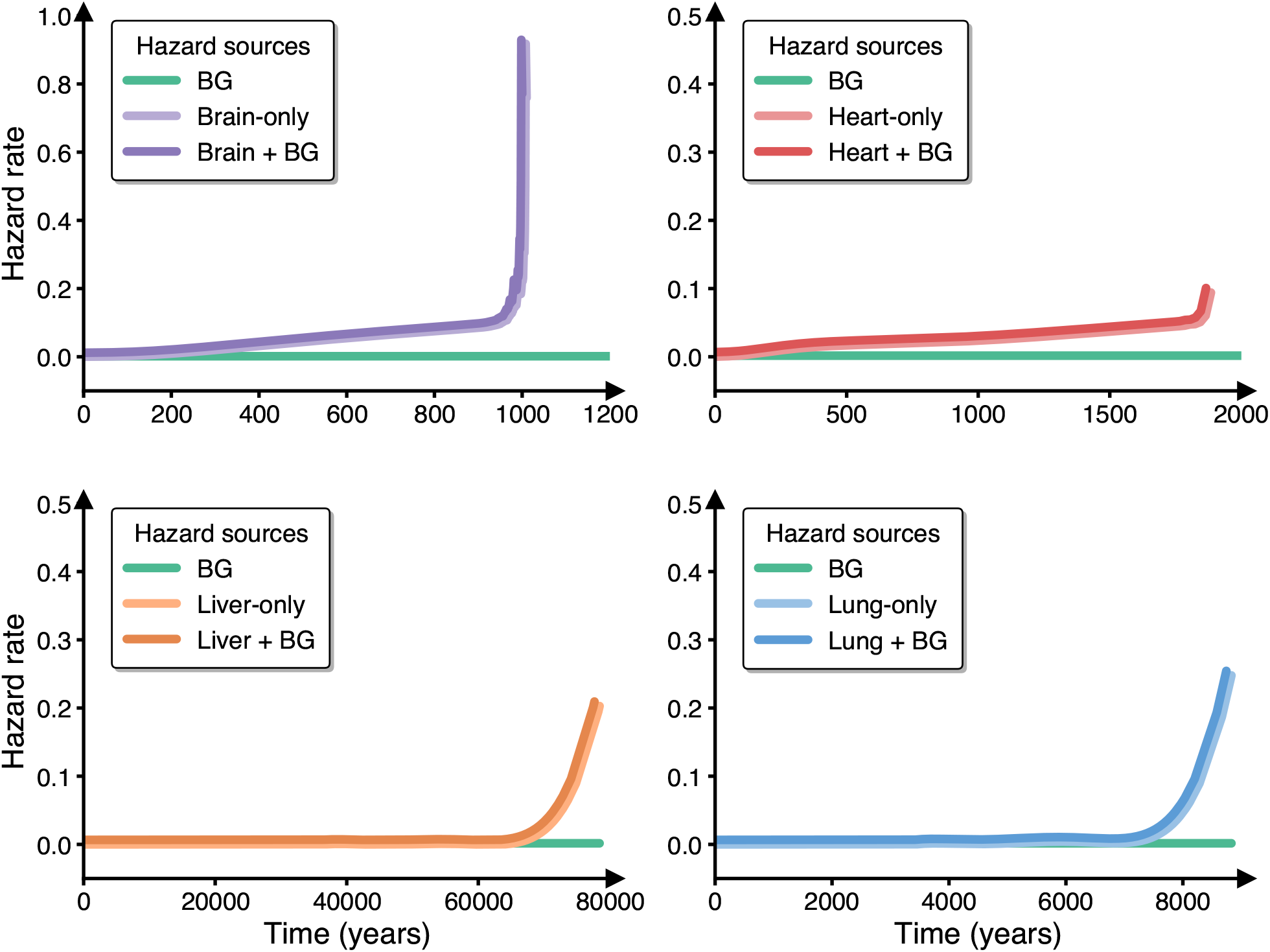
Hazard rates obtained for Model II (brain and heart), Model IIIA (liver), and Model IIIC (lung). Model IIIB is not shown, because it yielded constant hazard rate. Organ-only and organ+BG curves overlap each other and are shown with a slight shift from each other for visualization purposes.

**Extended Data Figure 5.**
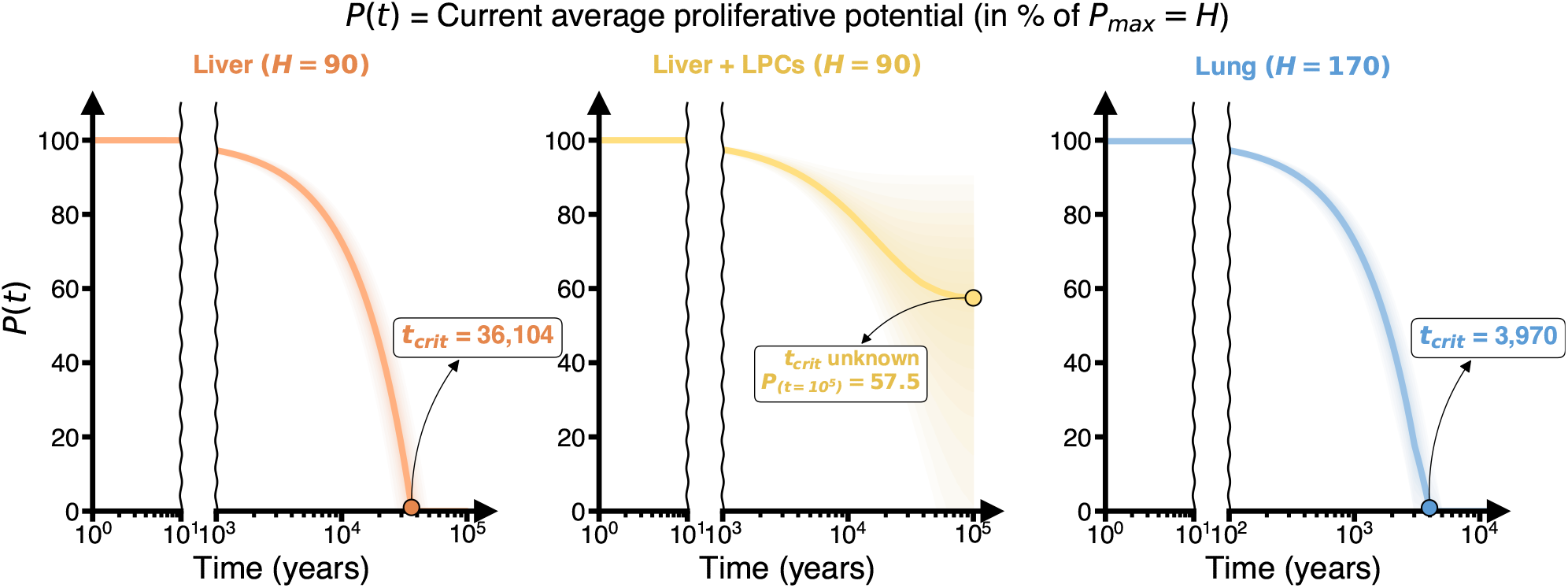
Proliferative potential dynamics for Model IIIA (hepatocytes only), Model IIIB (hepatocytes in the presence of LPC population), and Model IIIC (lung BBCs). Shaded regions represent percentile bands (from the 2.5th to the 97.5th in 5% increments) for the simulated *P*(*t*) trajectories; bright solid line corresponds to the median trajectory; the time at which the median potential trajectory crosses zero is annotated as *t*_crit_.

**Extended Data Figure 6.**
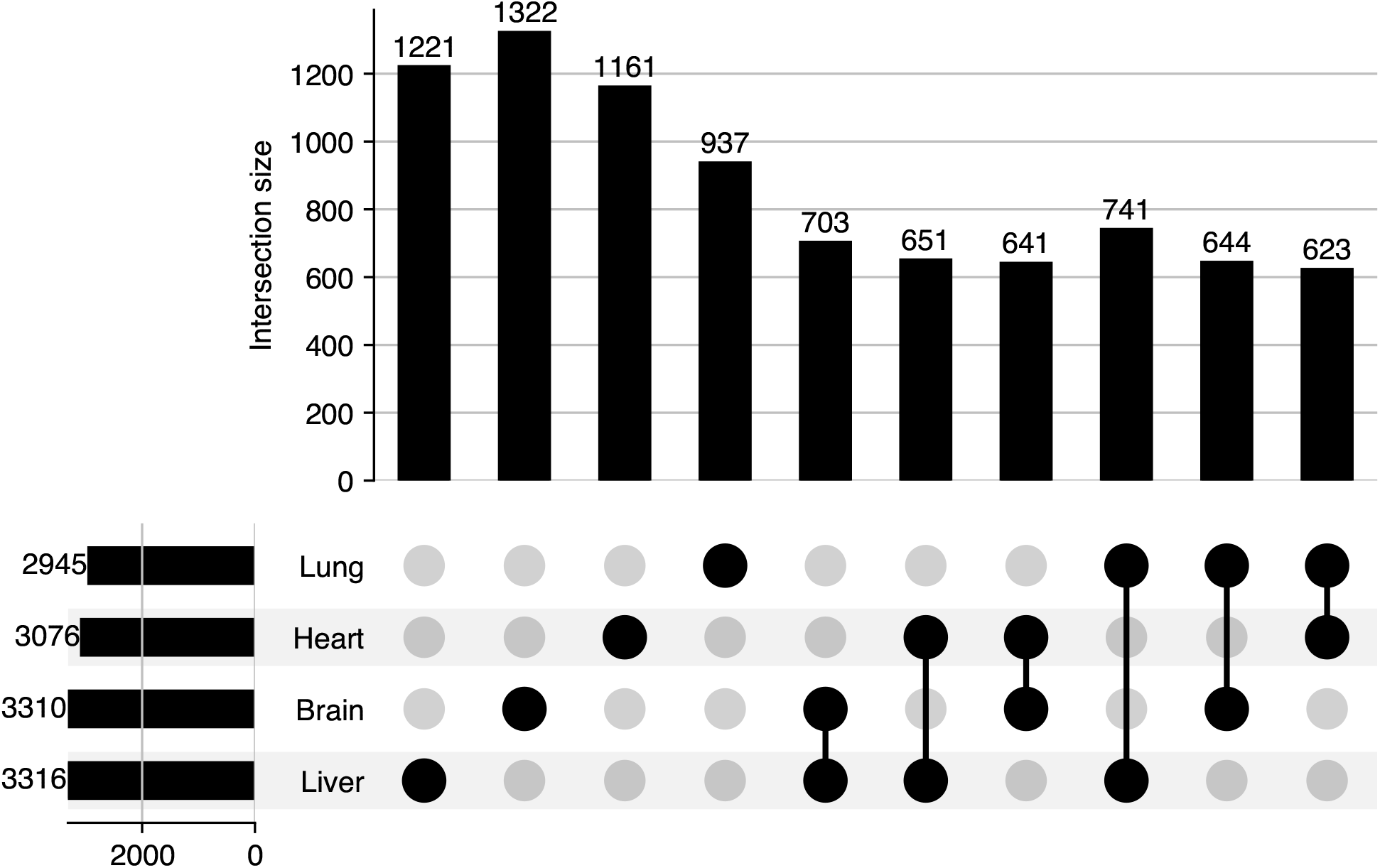
Essential gene signature sizes and their overlaps inferred for each studied organ.

**Extended Data Figure 7.**
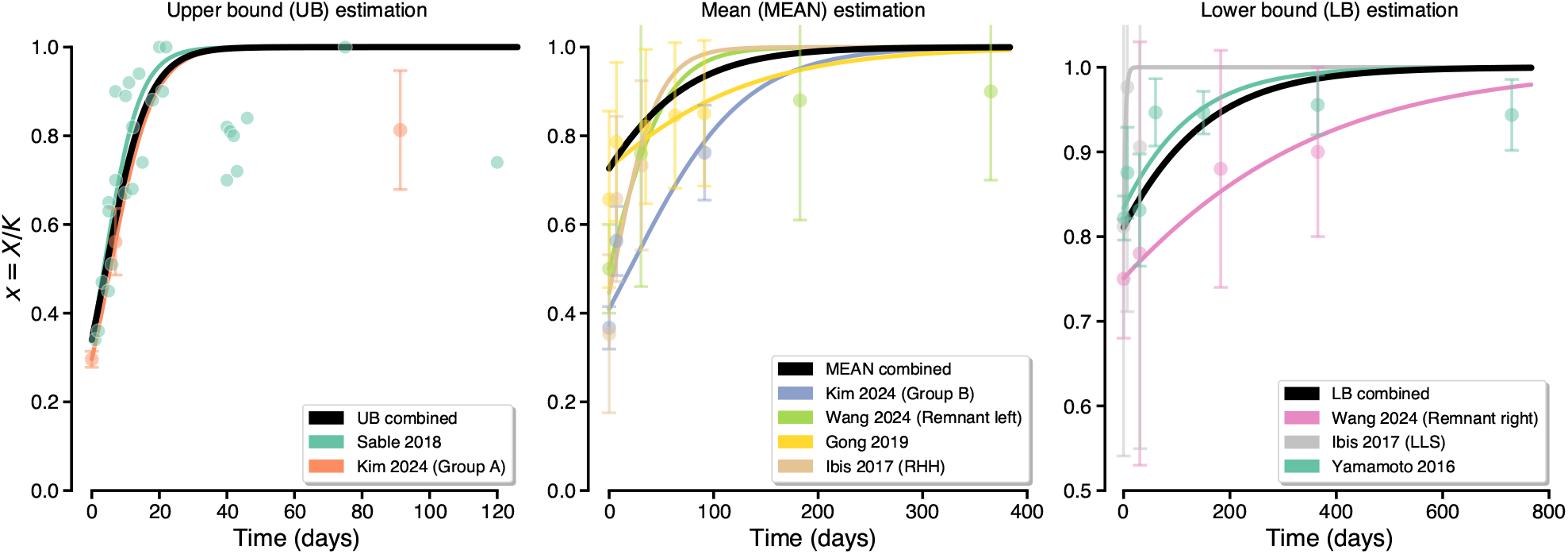
Hepatocyte *r* estimation from studies on partial hepatectomy at different remnant liver volumes *x*. For the Sable 2018 dataset, points represent individual *x* measurements. For all other datasets, points represent median values aggregated over a 1-day time period, and error bars represent 95% CI. RHH, right hemi-hepatectomy; LLS, left lateral sectionectomy.

**Extended Data Figure 8.**
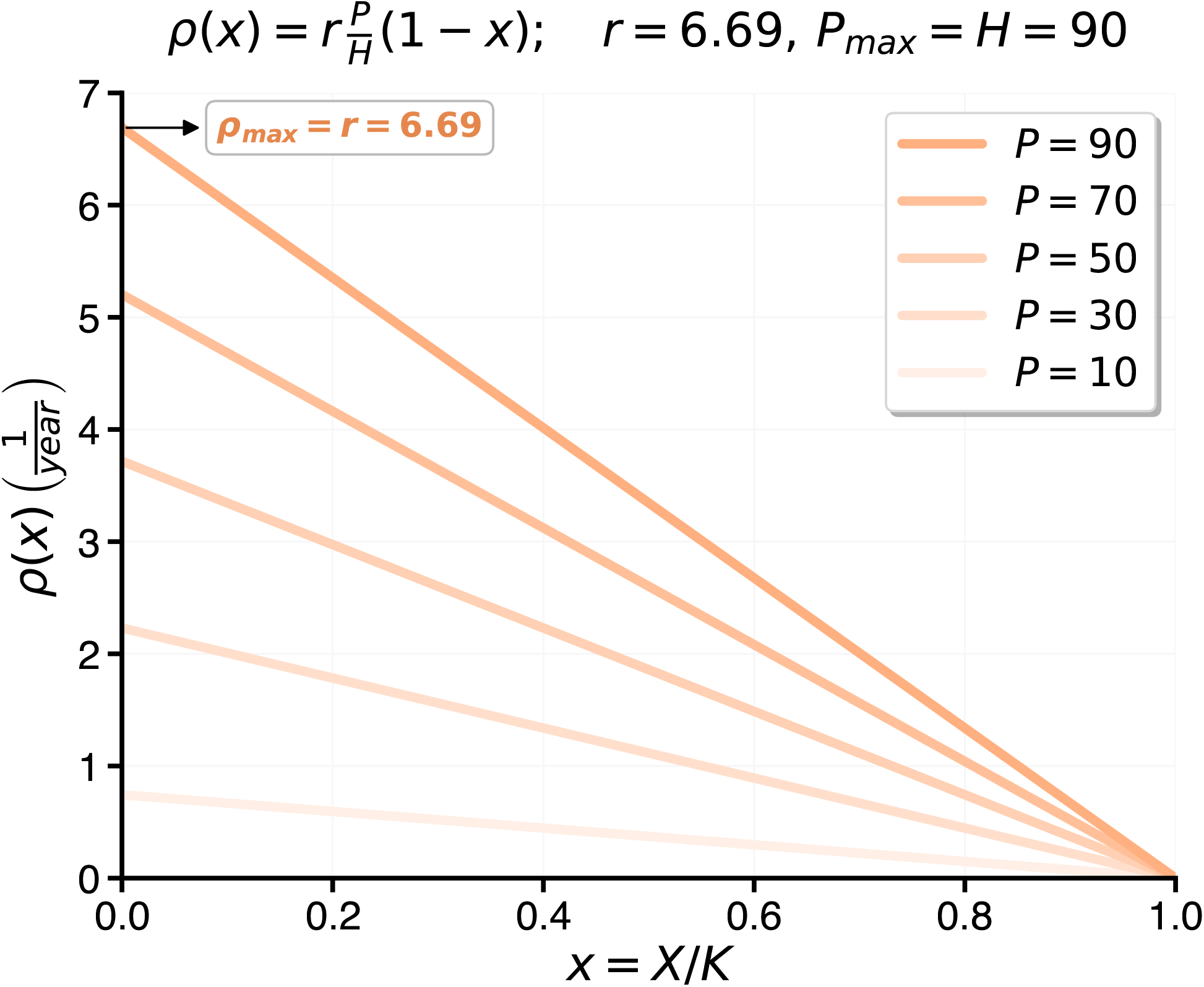
Hepatocyte replication rate *ρ* as a function of both current hepatocyte population *x* = *X/K* and average remaining proliferative potential *P*.

**Extended Data Figure 9.**
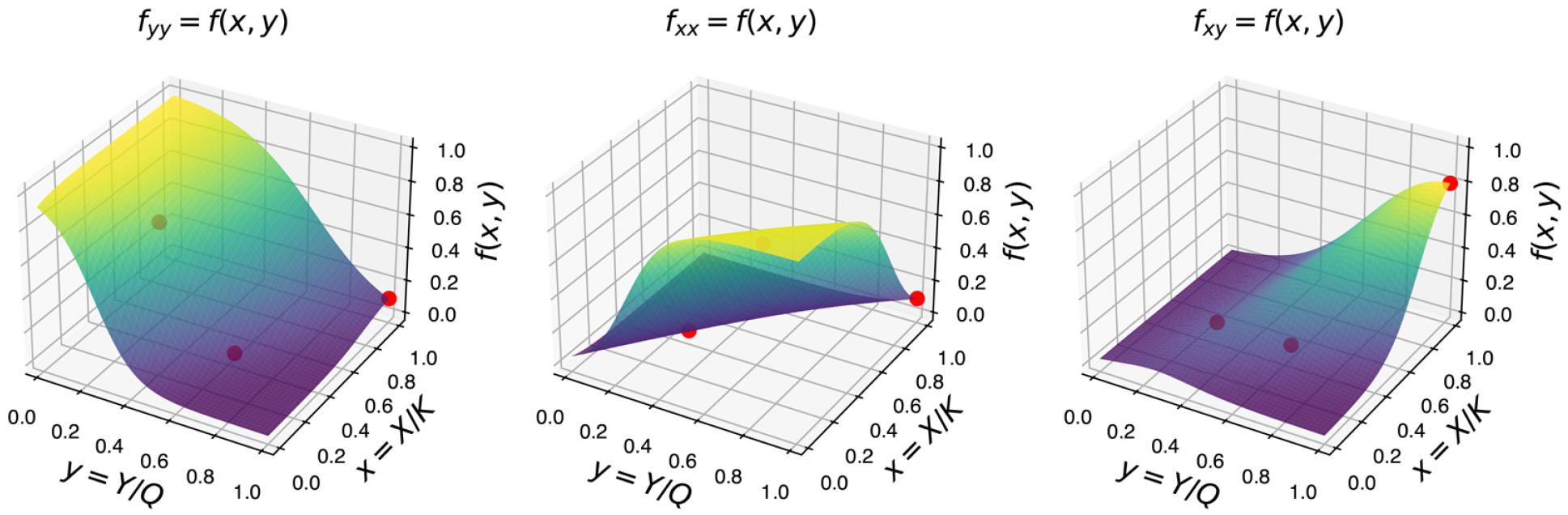
Probability landscapes of liver progenitor cell (LPC) division outcomes as functions of somatic cell (*x* = *X/K*) and LPC (*y* = *Y/Q*) population ratios. Each surface *f*_…_ = *f* (*x, y*) represents the probability that a stem cell undergoes symmetric self-renewal (*f*_*yy*_, left), symmetric differentiation (*f*_*xx*_, middle), or asymmetric division (*f*_*xy*_, right), with red markers indicating reference points selected for function fitting.

**Extended Data Figure 10.**
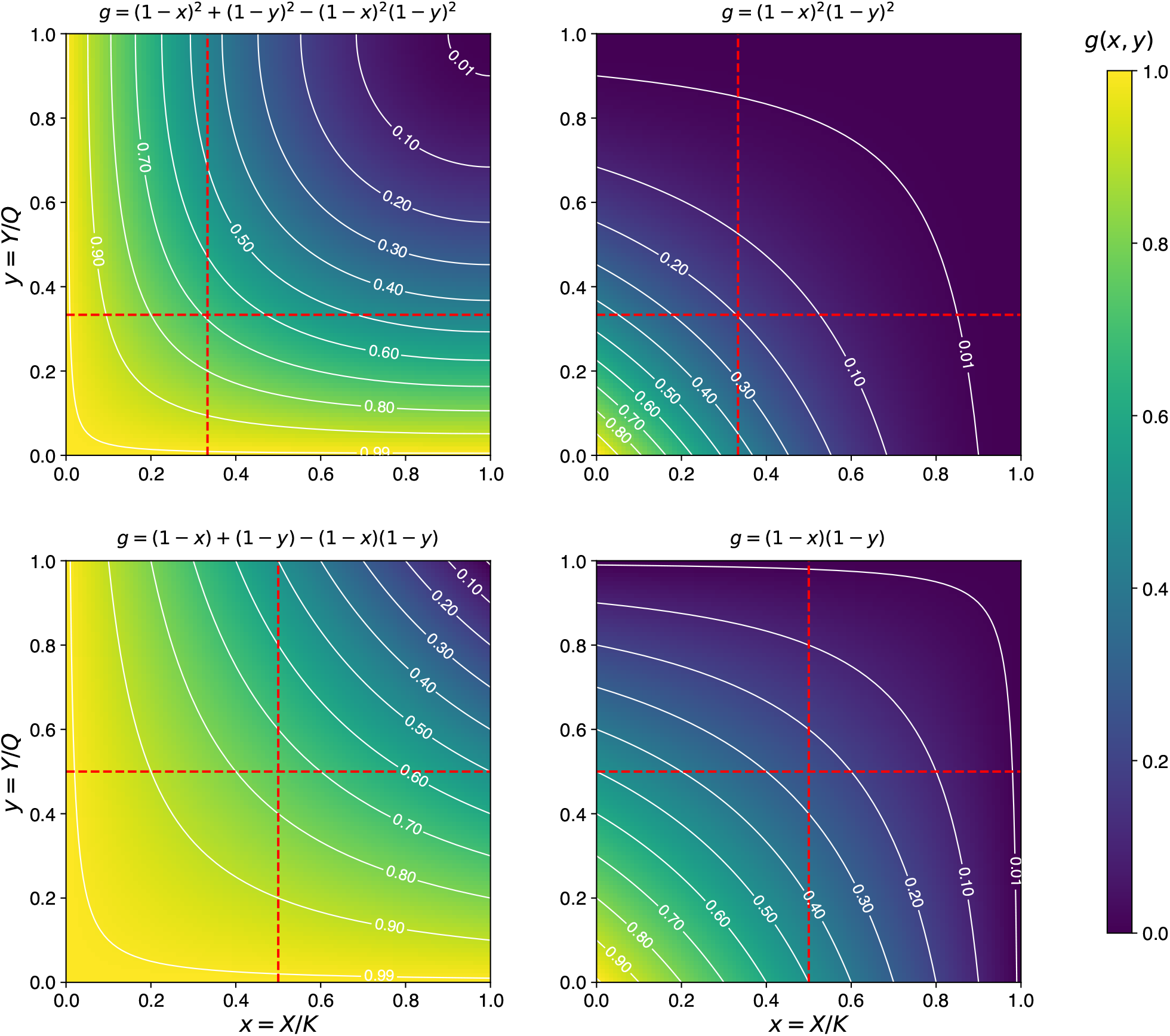
Different candidate gate functions *g* for stem cell renewal rate. Each gate accounts both for somatic *x* and stem cell *y* populations at a given time. Color bar corresponds to different *g* values. White countour lines represent specific *g* values from 0.01 to 0.99 in 0.1 increments. Dashed red lines represent *x* and *y* values at inflection points 1/3 (top left and top right) and 1/2 (bottom left and bottom right).

**Extended Data Figure 11.**
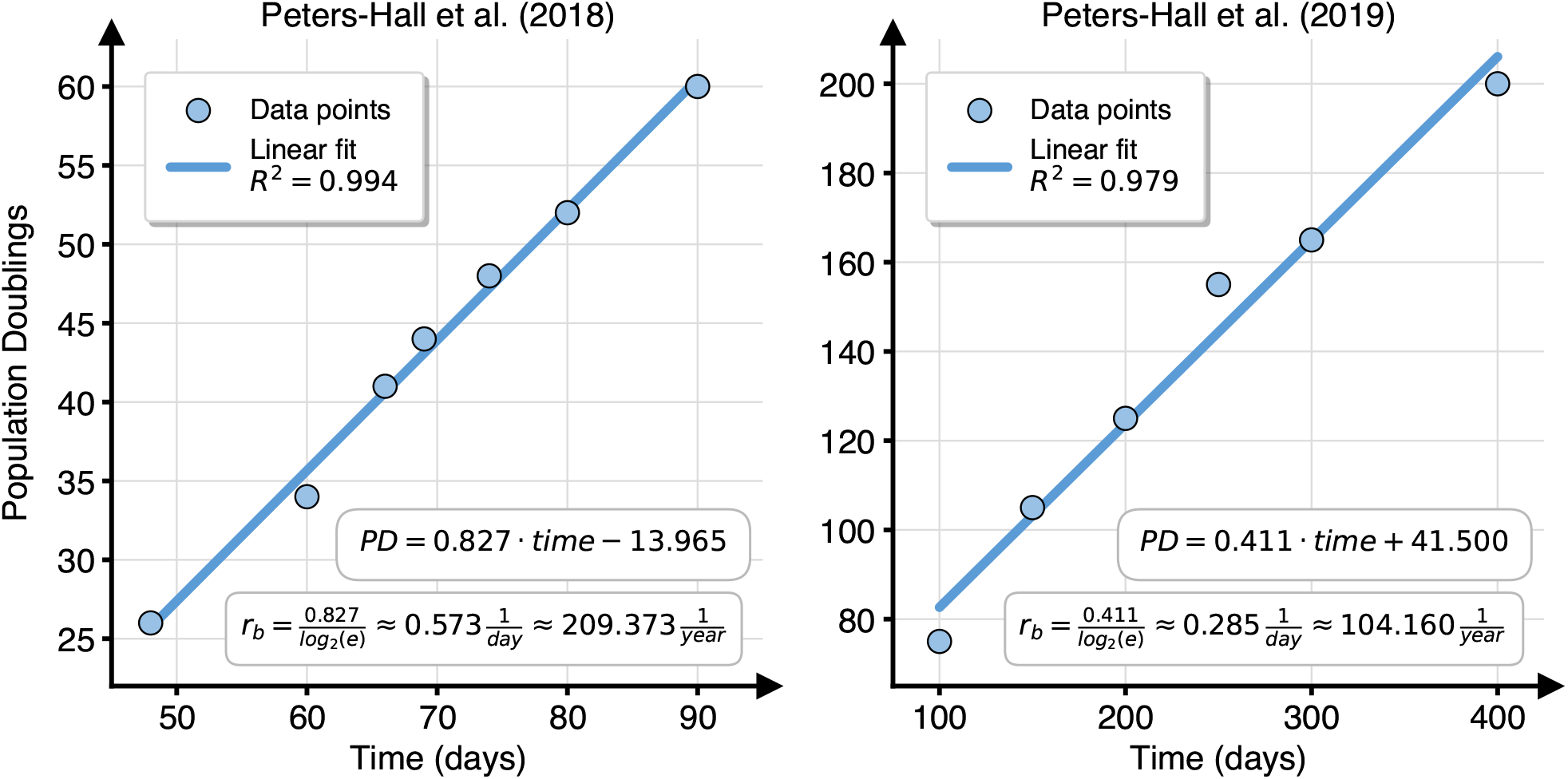
Bronchial basal cells (BBC) *r*_*b*_ estimation from studies on *in vitro* BBC expansion. Only the period of linear growth is shown. PD, population doublings.

**Extended Data Figure 12.**
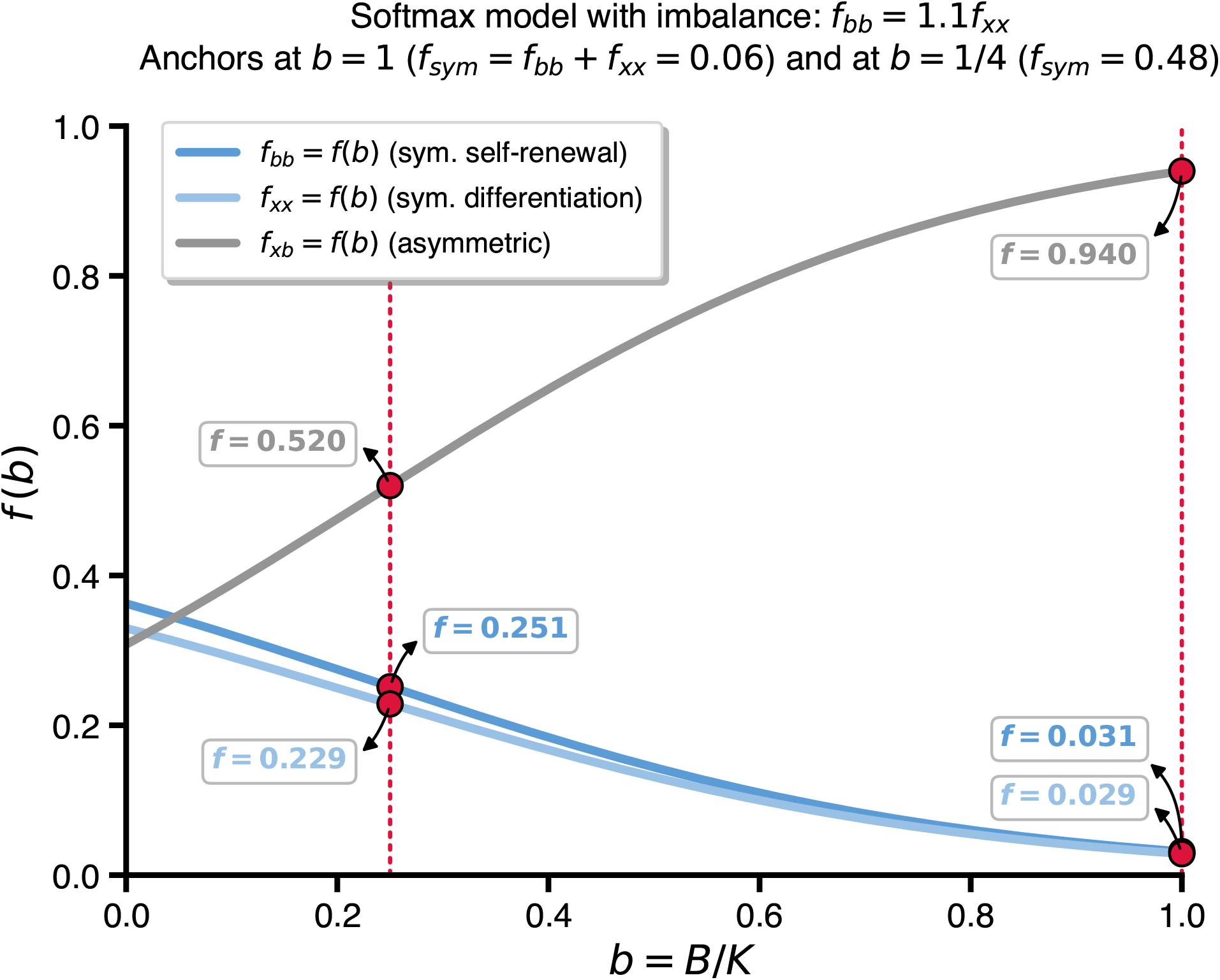
Probability functions of different BBC division outcomes as functions of BBC population ratio *b* = *B/K*. Each *f* (*b*) line represents the probability that a basal cell undergoes symmetric self-renewal (*f*_*bb*_), symmetric differentiation (*f*_*xx*_), or asymmetric division (*f*_*xb*_), with red markers indicating reference points selected for function fitting.

